# Spatiotemporal sequence of mesoderm and endoderm lineage segregation during mouse gastrulation

**DOI:** 10.1101/2020.06.09.142265

**Authors:** Simone Probst, Sagar, Jelena Tosic, Carsten Schwan, Dominic Grün, Sebastian J. Arnold

## Abstract

Anterior mesoderm (AM) and definitive endoderm (DE) progenitors represent the earliest embryonic cell types that are specified during germ layer formation at the primitive streak (PS) of the mouse embryo. Genetic experiments indicate that both lineages segregate from *Eomes* expressing progenitors in response to different NODAL signaling levels. However, the precise spatiotemporal pattern of the emergence of these cell types and molecular details of lineage segregation remain unexplored. We combined genetic fate labeling and imaging approaches with scRNA-seq to follow the transcriptional identities and define lineage trajectories of *Eomes* dependent cell types. All cells moving through the PS during the first day of gastrulation express *Eomes*. AM and DE specification occurs before cells leave the PS from discrete progenitor populations that are generated in distinct spatiotemporal patterns. Importantly, we don’t find evidence for the existence of progenitors that co-express markers of both cell lineages suggesting an immediate and complete separation of AM and DE lineages.

**Summary statement:** Cells lineages are specified in the mouse embryo already within the primitive streak where *Mesp1*+ mesoderm and *Foxa2*+ endoderm are generated in a spatial and temporal sequence from unbiased progenitors.

## Introduction

During mammalian gastrulation the pluripotent cells of the epiblast become lineage specified and form the three primary germ layers definitive endoderm (DE), mesoderm and (neuro-) ectoderm. Mesoderm and DE are generated at the posterior side of the embryo under the influence of elevated levels of the instructive signals of TGFß/NODAL, WNT and FGF. These signals induce an epithelial-to-mesenchymal transition (EMT) of epiblast cells at the primitive streak (PS) leading to their delamination and the formation of the mesoderm and DE cell layer. The nascent mesoderm layer rapidly extends towards the anterior embryonic pole by cell migration between the epiblast and the visceral endoderm (VE) (reviewed by (Arnold and Robertson 2009; Rivera-Pérez et al. 2003)). DE progenitors migrate from the epiblast together with mesoderm cells, before they eventually egress into the VE layer to constitute the DE (reviewed by (Rivera-Pérez and Hadjantonakis 2014; Viotti, Foley, et al. 2014)).

Current concepts suggest that different cell fates are specified according to the time and position of cell ingression through the PS reflecting different instructive signaling environments (Rivera-Pérez and Hadjantonakis 2014). However, the precise morphogenetic mechanisms guiding the emergence of various cell types along the PS still remain uncertain. This is at least in parts due to the lack of detailed knowledge about the precise timing and location of individual cells becoming lineage specified, and the challenge to exactly determine the signaling pathway activities during fate commitment.

Clonal cell labeling and transplantation experiments have proposed the gross patterns and dynamics of cell specification during gastrulation, which have been represented in fate maps of the epiblast and the early germ layers (Tam and Behringer 1997; Lawson 1999). Accordingly, first mesoderm cells delaminate from the newly formed PS at the proximal posterior pole of the embryo and give rise to extraembryonic mesoderm cells (ExM). These migrate proximally and anteriorly to contribute to the mesodermal components of the amnion, chorion, and the yolk sac (Parameswaran and Tam 1995; Kinder et al. 1999). Embryonic anterior mesoderm (AM) giving rise to cardiac and cranial mesoderm follows shortly after ExM (Kinder et al. 1999). As the PS elongates towards the distal embryonic pole other mesoderm subtypes and DE are generated. The distal domain of the PS (referred to as anterior PS, APS) generates DE and axial mesoderm progenitors, giving rise to the node, notochord and prechordal plate mesoderm (Kinder et al. 2001; Lawson et al. 1991). Additional mesoderm subtypes, such as lateral plate, paraxial and intermediate mesoderm are generated between the APS and the proximal PS (Lawson et al. 1991; Kinder et al. 1999; Tam et al. 1997; Parameswaran and Tam 1995).

TGFß/NODAL and WNT signals are indispensable for gastrulation onset (Brennan et al. 2001; Conlon et al. 1994; Liu et al. 1999) and genetic experiments revealed that graded levels of NODAL and WNT signaling instruct distinct lineage identities during gastrulation (Vincent et al. 2003, Dunn et al., 2004; reviewed by (Robertson 2014; Arkell et al. 2013)). The T-box transcription factor *Eomes* is a transcriptional target of NODAL/SMAD2/3 signaling (Brennan et al. 2001; Teo et al. 2011; Kartikasari et al. 2013) and is crucial for the specification of all DE and AM progenitors ((Arnold et al. 2008; Costello et al. 2011; Probst and Arnold 2017). Another T-box transcription factor, *Brachyury* (T) is essential for the formation of posterior mesoderm starting from E7.5. Thus, the specification of all types of mesoderm and endoderm (ME) relies on either of the two T-box factors *Eomes* or *Brachyury* (Tosic et al. 2019). Experiments using differentiating human embryonic stem cells showed that EOMES directly binds and regulates the expression of DE genes together with SMAD2/3 (Teo et al. 2011). Similarly in the mouse embryo DE specification relies on high NODAL/SMAD2/3 signaling levels (Dunn et al. 2004; Vincent et al. 2003). In contrast, in the presence of low or even absent NODAL/SMAD2/3 signals, EOMES activates transcription of key determinants for anterior mesoderm, including *Mesp1* (Saga et al. 1999; Lescroart et al. 2014; Kitajima et al. 2000; Costello et al. 2011; van den Ameele et al. 2012).

Recently single cell RNA sequencing (scRNA-seq) analyses allowed for a more detailed view on the cellular composition of embryos during gastrulation stages including the identification of previously unknown rare and transient cell types (Scialdone et al. 2016; Mohammed et al. 2017; Wen et al. 2017; Lescroart et al. 2018; Pijuan-Sala et al. 2019). Despite the insights into the molecular mechanisms of cell lineage specification, questions about the emergence of the two *Eomes* dependent cell lineages, AM and DE, remain unresolved. It is still unclear if both cell populations are generated simultaneously from a common progenitor, and when and where lineage separation occurs. Answers to these questions are required for a comprehensive view on how suggested differences in the signaling environment impact on lineage specification of mesoderm and DE identities that are generated in close proximity within the epiblast of early gastrulation stage embryos that consist of only a few hundred cells (Snow 1977).

In this study we used embryo imaging and genetic fate mapping approaches by novel reporter alleles in combination with molecular characterization by scRNA-seq to delineate the spatiotemporal patterns of *Eomes* dependent lineage specification. We show that AM and DE progenitors segregate already within the PS into distinct cell lineages. AM progenitors leave the PS earlier and at more proximal regions than DE, demonstrating a clear spatial and temporal separation of lineage specification. The analysis of scRNA-seq experiments suggest that AM and DE progenitors are immediately fate segregated and *Eomes* positive progenitors that co-expresses DE and AM markers were not found. This suggests that bipotential *Eomes* positive progenitors rapidly progress into either AM or DE lineage specified cell types preceding cell ingression at the PS.

## Results

### Eomes marks all cells leaving the PS during the first day of gastrulation

We used our previously described *Eomes*^*mTnG*^ fluorescent reporter allele to observe the emergence of *Eomes* dependent cell lineages during gastrulation. This reporter allele labels *Eomes* positive cells with membrane bound Tomato (mT) and nuclear GFP (nG) ((Probst et al. 2017) and Fig. 1A-F). Embryos at stages shortly preceding gastrulation onset (E6.25) show labeling within the cells of the posterior epiblast before cells leave the PS (Fig. 1A). *Eomes* positive cells are also detected in the VE where it has functions in AVE induction (Nowotschin et al. 2013). Reporter expression in the epiblast persists until E7.5 marking the prospective cells of the AM and DE (Fig. 1B-D). Importantly, all cells leaving the PS are EOMES positive during these early gastrulation stages suggesting a general requirement of *Eomes* for the specification of early lineages (Fig. 1B-E, G, H). Accordingly, the maximum intensity projection (MIP) of z-stacks at E7.5 shows that the endoderm layer, that at this stage mainly consists of DE cells, is composed of *Eomes* reporter positive cells (Fig. 1F). In the endoderm layer a few reporter negative cells can be detected, which most likely represent VE cells, that loose reporter expression at around E7.25 (Fig. 1C, F). Since the fluorescent reporter proteins are more stable than the endogenous protein (Probst et al. 2017), we additionally performed immunofluorescence (IF) staining for EOMES at E7.25 and E7.5, showing the presence of EOMES protein in the cells of the posterior epiblast and in cells of the mesoderm and endoderm layers (Fig. 1G, H). *Eomes* mRNA expression is rapidly downregulated at E7.5 (Ciruna and Rossant 1999), and EOMES protein is undetectable about 12 hours later (Probst et al. 2017). In conclusion, mesoderm and endoderm progenitors generated during the first day of gastrulation from E6.5 to E7.5 are exclusively descendants of *Eomes* expressing cells (Fig. 1P). These constitute the progenitors of AM and DE as previously shown by *Eomes*^Cre^-mediated fate-labeling (Costello et al. 2011). This suggests that other mesodermal lineages, which are *Eomes* independent, leave the PS at later timepoints after the downregulation of *Eomes* expression in the PS.

**Figure 1:**
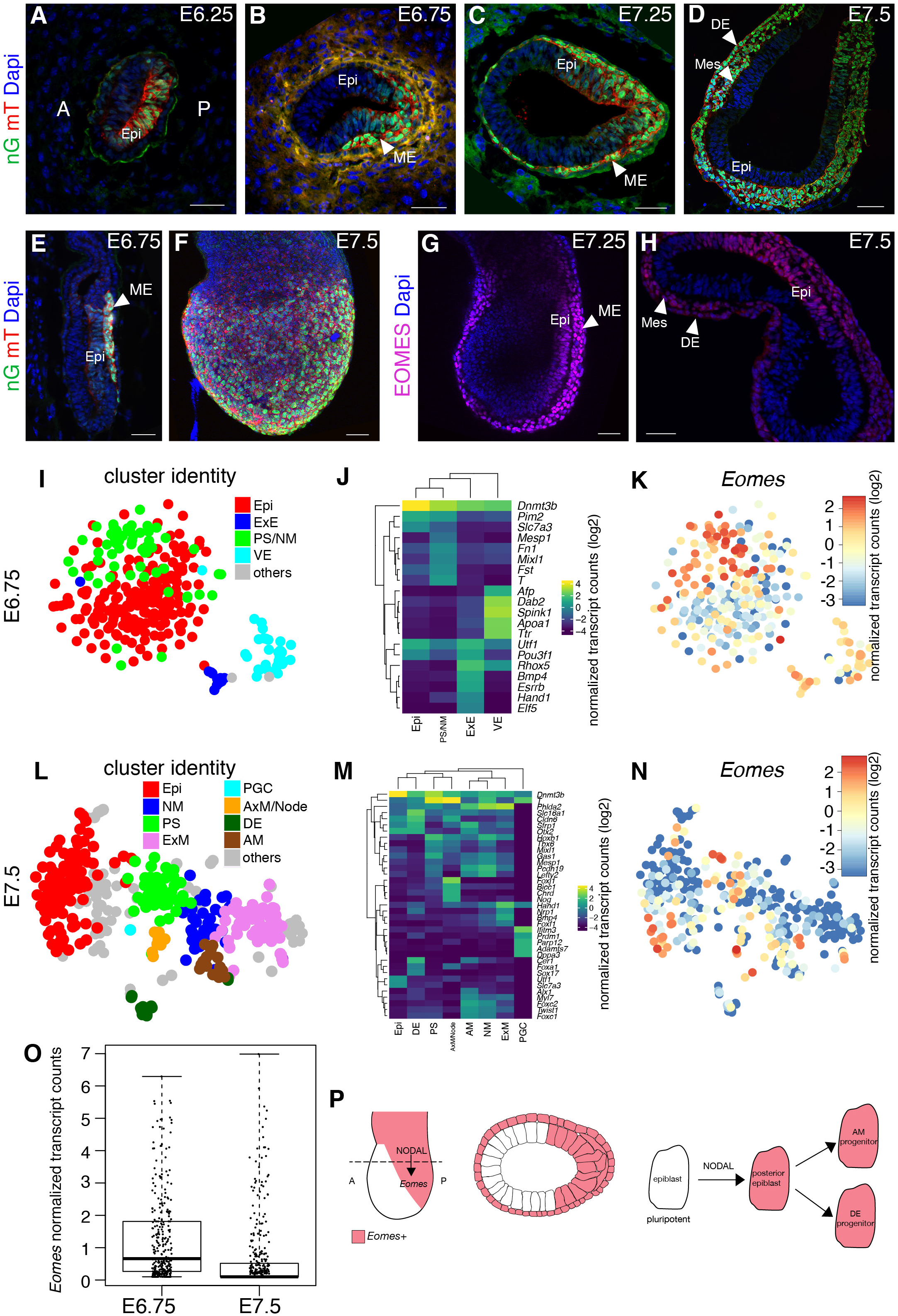
All cells of the posterior epiblast and AM and DE progenitors between E6.25 and E7.5 express *Eomes*. **A-H**) Immunofluorescence (IF) staining of *Eomes*^*mTnG*^ and wild type (wt) embryos (n ≥ 3 embryos). In this and all following figures embryos are oriented with anterior (A) to the left and posterior (P) to the right. **A-D**) Transverse sections of *Eomes*^*mTnG*^ embryos showing nuclear GFP (nG) and membrane bound Tomato (mT) in *Eomes* expressing cells. **A**) Before gastrulation onset at E6.25, *Eomes*^*mTnG*^ positive cells mark the posterior half of the proximal epiblast (Epi) and thus the prospective site of primitive streak formation. **B**) At E6.75 and (**C**) E7.25 mesoderm and endoderm (ME) cells have ingressed through the primitive streak and migrate towards the anterior side of the embryo. The posterior Epi and all ME cells are positive for *Eomes*^*mTnG*^. **D**) At E7.5 the posterior epiblast remains *Eomes*^*mTnG*^ positive, but in a restricted domain compared to earlier stages. The *Eomes*^*mTnG*^ positive mesoderm (Mes) wings have migrated to the anterior and *Eomes*^*mTnG*^ positive DE cells integrated into the outer endoderm layer. **E**) Sagittal section of an E6.75 embryo showing *Eomes*^*mTnG*^ expression in the nascent mesoderm layer. **F**) Maximum intensity projection (MIP) of an E7.5 *Eomes*^*mTnG*^ embryo. **G**) Sagittal section of an E7.25 wt embryo stained for EOMES showing remainng expression in the Epi and ME (arrowhead). **H**) Transverse section of an E7.5 wt embryo stained for EOMES. Endogenous EOMES protein remains present in the posterior Epi and in the ME layer (arrowhead); however, protein levels are reduced in the more anterior mesoderm and DE. Scale bars 50 μm. **I-O**) scRNA-seq of wt embryos at E6.75 and E7.5. **I, L**) t-SNE plots with assigned identities to different clusters at (**I**) E6.75 and (**L**) E7.5. Anterior mesoderm (AM), axial mesoderm (AxM), definitive endoderm (DE), epiblast (Epi), nascent mesoderm (NM), extraembryonic ectoderm (ExE), extraembryonic mesoderm (ExM), primordial germ cell (PGC), primitive streak (PS), and visceral endoderm (VE). **J, M**) Heat maps of selected marker genes for the clusters indicated in (**I, L**) at E6.75 (**J**) and at E7.5 (**M**). Scale bar represents log2 normalized transcript counts. **K, N**) t-SNE plots showing the expression of *Eomes* in single cells at E6.75 (**K**) and E7.5 (**N**), the scale represents log2 normalized transcript counts. **O**) Box plot showing the expression levels of *Eomes* by normalized transcript counts in single cells at both timepoints indicating a higher proportion of *Eomes* expressing cells at E6.75. **P**) Schematic illustrating the generation of *Eomes* dependent cell lineages in the posterior embryo at E6.75. Nodal is required for the induction of *Eomes* in the posterior half of the epiblast to induce the early AM and DE progenitors.

To molecularly characterize the *Eomes* dependent cell types during early gastrulation we performed scRNA-seq of cells collected from E6.75 and E7.5 embryos (Fig. 1I-O). 289 handpicked cells from 14 E6.75 embryos and 371 cells isolated by FACS from E7.5 pooled litters were included in the scRNA-seq analysis. To find transient progenitor populations within the epiblast we clustered the cells using RaceID3 (Herman et al. 2018), an algorithm specifically developed for the identification of rare cell types within scRNA-seq data (Grün et al. 2015). This analysis defined seven different cell clusters at E6.75 and 18 clusters at E7.5 (Fig. S1A, B).

The tissue identities of clusters were assigned by the presence of differentially upregulated marker genes in each cluster compared to the rest of the cells (Fig. 1I, J, L, and M, and Tables S1, S2). The heatmap representations indicate specifically expressed marker genes in different cell types (Fig. 1J, M). At E7.5 RaceID identified rare cells such as one single E7.5 primordial germ cell (PGC) (Fig. 1L, M and Table S2). No extraembryonic clusters were detected at E7.5, as the embryos were cut below the chorion during dissection excluding the ExE and VE cells are mixed with DE cells. The comparison of the t-SNE maps at E6.75 and E7.5 (Fig. 1I, L) shows that at E6.75 epiblast and PS/ native mesoderm (NM) cells cluster closely to each other and only extraembryonic tissues (ExE and VE) form clearly separated clusters (Fig. 1I). In contrast, at E7.5 separable clusters can be detected within the embryonic cell clusters, demonstrating the increase in expression diversity of embryonic cell types between E6.75 to E7.5 (Fig. 1L). Of note, no specific subclusters could be found within the epiblast cells at E6.75 which could represent trajectories towards mesoderm and endoderm progenitors.

Remarkably, at E6.75 *Eomes* is expressed in 209 of 289 cells (72% of all cells), showing highest expression in the PS/NM cluster (Fig. 1K, and Fig. S1C) and weaker expression in the epiblast cluster, which is in agreement with the immunofluorescence staining (Fig. 1B). In addition, *Eomes* expression is found in the extraembryonic clusters, ExE and VE (Fig. 1K and Fig. S1C). At E7.5 *Eomes* expression is still present in a subset of epiblast cells, the PS, the NM, the node and the mesoderm (AM and ExM) and DE clusters (Fig. 1N, Fig. S1D). However, only 35% of cells show RNA expression, whereas EOMES protein is still broadly detected (Fig. 1H, N). Thus, at E6.75 *Eomes mRNA* is generally expressed in more cells and at higher levels than at E7.5 (Fig. 1O). In summary, fluorescent reporter and scRNA-seq analyses demonstrate that *Eomes* marks prospective AM and DE progenitors within the posterior region of the epiblast from stages preceding the onset of cell ingression through the PS and until cells are present in the mesoderm and endoderm cell layer. ScRNA-seq analyses indicate that the embryonic *Eomes* positive cells at E6.75 are quite similar to each other and the only clusters identified were the PS and early mesoderm progenitors (PS/NM Fig. 1I) and intermediate progenitor clusters were not detected.

### A novel Mesp1^mVenus^ allele identifies Eomes dependent AM progenitors

*Mesp1* represents one of the earliest markers of mesoderm within the *Eomes* positive cell population and is a direct transcriptional target gene of EOMES (Costello et al. 2011). Lineage tracing with a *Mesp1-Cre* allele shows that it faithfully labels the ExM and the AM (Saga et al. 1999; Lescroart et al. 2014; Lescroart et al. 2018; S. S.-K. Chan et al. 2013). Thus, to distinguish mesoderm and DE progenitors during the first day of germ layer formation we generated a fluorescent *Mesp1*^*mVenus*^ reporter allele. We inserted the coding sequence of membrane bound Venus (mV) into the start codon of the *Mesp1* gene locus followed by a T2A cleavage site and the *Mesp1* coding sequence to maintain functional *Mesp1* expression from the resulting reporter allele (Fig. 2A-C). Homozygous *Mesp1*^*mVenus*^ (*Mesp1*^*mV*^) mice are viable and fertile, demonstrating sufficient *Mesp1* expression from the reporter allele.

**Figure 2:**
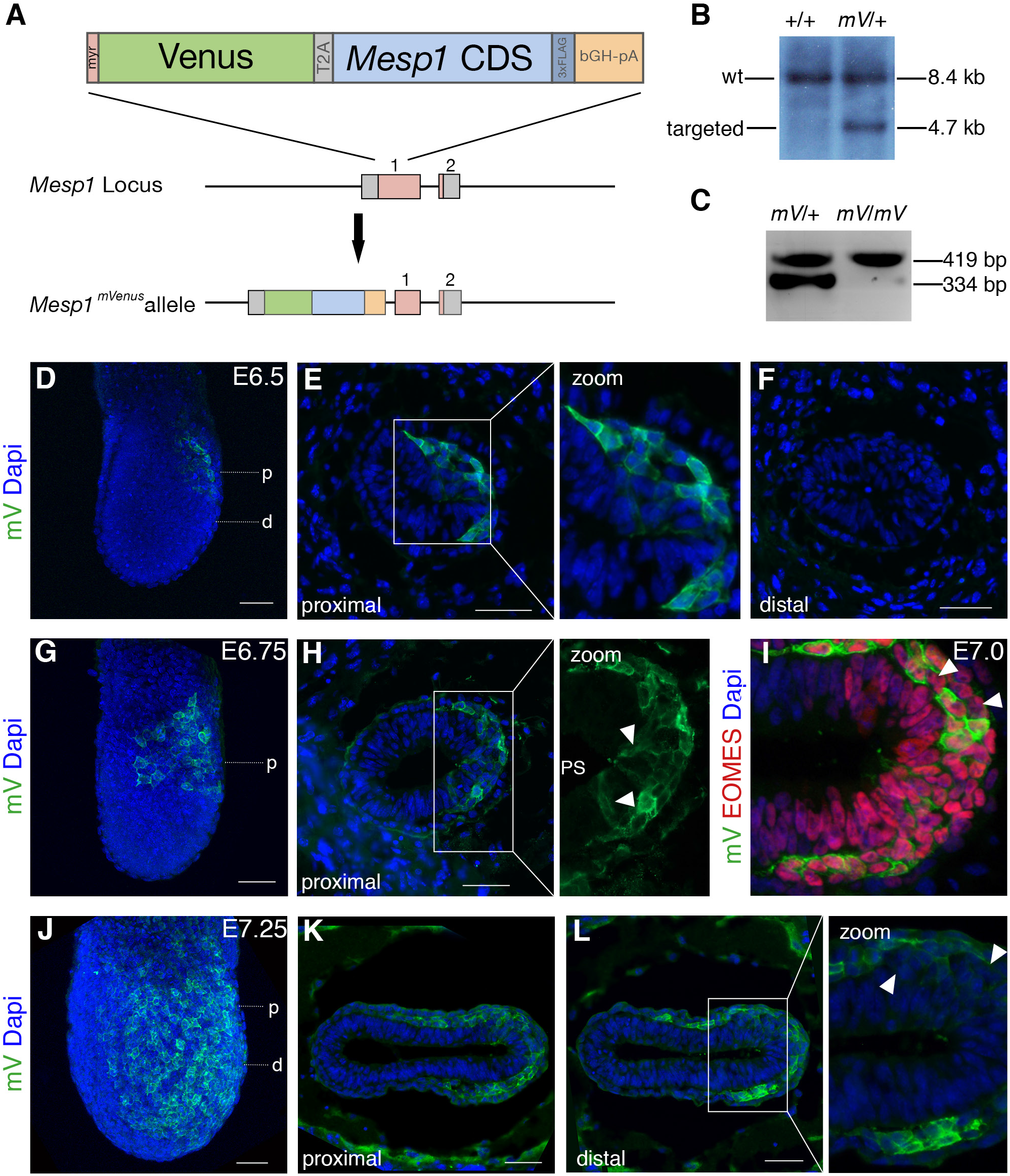
Generation of a novel *Mesp1*^*mVenus*^ allele to identify the *Eomes* dependent mesoderm progenitors. **A**) Schematic of the *Mesp1*^*mVenus*^ allele. The sequence for a membrane-targeted (myr) Venus protein (mV) and the *Mesp1* coding sequence (CDS) were inserted into the ATG of the *Mesp1* gene by homologous recombination to generate the *Mesp1*^*mVenus*^ (*Mesp1*^*mV*^) allele. **B**) Southern blot analysis of targeted ES cell clones showing a wt (+/+) and a correctly targeted (*mV*/+) clone. The wt band is detected at 8.4 kb and the targeted band is detected at 4.7. **C**) Genotyping PCR of a heterozygous (*mV*/+) and a homozygous (*mV/mV*) mouse showing wt band at 334 bp and the *mV* band at 419 bp. **D-L**) Immunofluorescence stainings with anti-GFP antibody to enhance mV protein in *Mesp1*^*mV*^ embryos (n ≥ 3 embryos). **D-F**) First *Mesp1*^*mV*^ positive cells appear during initiation of gastrulation at E6.5. **E, F**) Transverse sections at E6.5 show that early gastrulating cells in the proximal embryo are positive for *Mesp1*^*mV*^ expression (**E**), while in more distal regions the PS has not yet formed (**F**). **G**) By E6.75 *Mesp1*^*mV*^ positive cells rapidly migrate proximally towards the extraembryonic domain and anteriorly. **H**) *Mesp1*^*mV*^ positive cells are already detected in the epithelial PS (arrowheads in zoom). **I-L**) By E7.0 and E7.25 the *Mesp1*^*mV*^ positive cells constitute a large population within the mesoderm layer. **I**) Costaining with anti-EOMES antibody shows that *Mesp1*^*mV*^ expressing cells represent a subpopulation of EOMES positive cells (arrowheads indicate few *Mesp1*^*mV*^ negative cells). **J, K**) At E7.25 the proximal mesoderm layer contains mainly *Mesp1*^*mV*^ positive cells and the PS is also positive for *Mesp1*^*mV*^. **L**) More distally *Mesp1*^*mV*^ positive cells are intermixed with *Mesp1*^*mV*^ negative cells and the PS contains no *Mesp1*^*mV*^ cells. The scale bars represent 50 μm. **D**, **G**, **J**, show MIPs. The approximate levels of the transverse sections are indicated in the MIPs. p, proximal; d, distal.

We analyzed the emergence of earliest *Eomes* dependent mesoderm progenitors in *Mesp1*^*mV*^ embryos and found first *Mesp1* positive cells already at E6.5 in the proximal epiblast during early PS formation (Fig. 2D, E). At this stage the PS has not yet extended towards the distal part of the embryo, and also no *Mesp1*^*mV*^ positive cells are present in distal portions of the epiblast (Fig. 2F). Notably, almost all cells leaving the early proximal PS show *Mesp1* reporter expression identifying them as mesoderm progenitors (Fig. 2E, zoom). Once *Mesp1*^*mV*^ positive cells leave the PS they rapidly migrate proximally and anteriorly to their destinations of ExM and AM (Fig. 2G). Importantly, *Mesp1*^*mV*^ positive cells are already detected in the epithelial portion of the PS (Fig. 2H zoom, arrowheads), indicating that mesoderm fate specification takes place before cells delaminate from the epiblast. At E7.25 the mesodermal wings have migrated far anteriorly (Fig. 2J). *Mesp1*^*mV*^ positive cells constitute the major population within the EOMES positive mesodermal layer, (Fig. 2I). While the proximal mesoderm is mainly composed of *Mesp1*^*mV*^ positive cells, increasing numbers of *Mesp1*^*mV*^ negative cells were found more distally (Fig. 2I, L, zoom, arrowheads). At E7.25 nascent *Mesp1*^*mV*^ cells are still emerging from the proximal PS (Fig. 2K), while more distally no *Mesp1*^*mV*^ positive cells were detected in the PS (Fig. 2L). *Mesp1* therefore marks the earliest population of mesoderm progenitors that are continuously produced between E6.5 and E7.5 from *Eomes* expressing cells. *Mesp1* positive progenitors are present throughout the mesoderm layer but they are preferentially generated in the proximal domains of the PS.

### DE and AM progenitors become fate specified in different regions of the epiblast

Next, we investigated the spatial distribution of the *Eomes* dependent cell lineages (Costello et al. 2011; Arnold et al. 2008). To simultaneously detect DE and AM progenitors we used FOXA2 immunofluorescence staining of embryos carrying the *Mesp*^*mV*^ reporter allele. Previous reports and our data show that *Foxa2* is expressed in the VE and during gastrulation from E6.5 onwards in the epiblast, the APS/node, and its derivatives DE and axial mesoderm (Fig. 3, (Ang et al. 1993; Sasaki and Hogan 1993; Monaghan et al. 1993; Viotti, Nowotschin, et al. 2014)). Previous lineage tracing by Cre-induced recombination and imaging by fluorescent reporters showed that *Foxa2* expression faithfully labels DE progenitors (Park et al. 2008; Frank et al. 2007; Imuta et al. 2013). While *Foxa2* is not strictly required for the initial egression of DE cells into the VE layer (Viotti, Nowotschin, et al. 2014), *Foxa2* deficient embryos lack foregut and midgut DE formation and only generate hindgut endoderm (Dufort et al. 1998; Ang and Rossant 1994; Weinstein et al. 1994).

**Figure 3:**
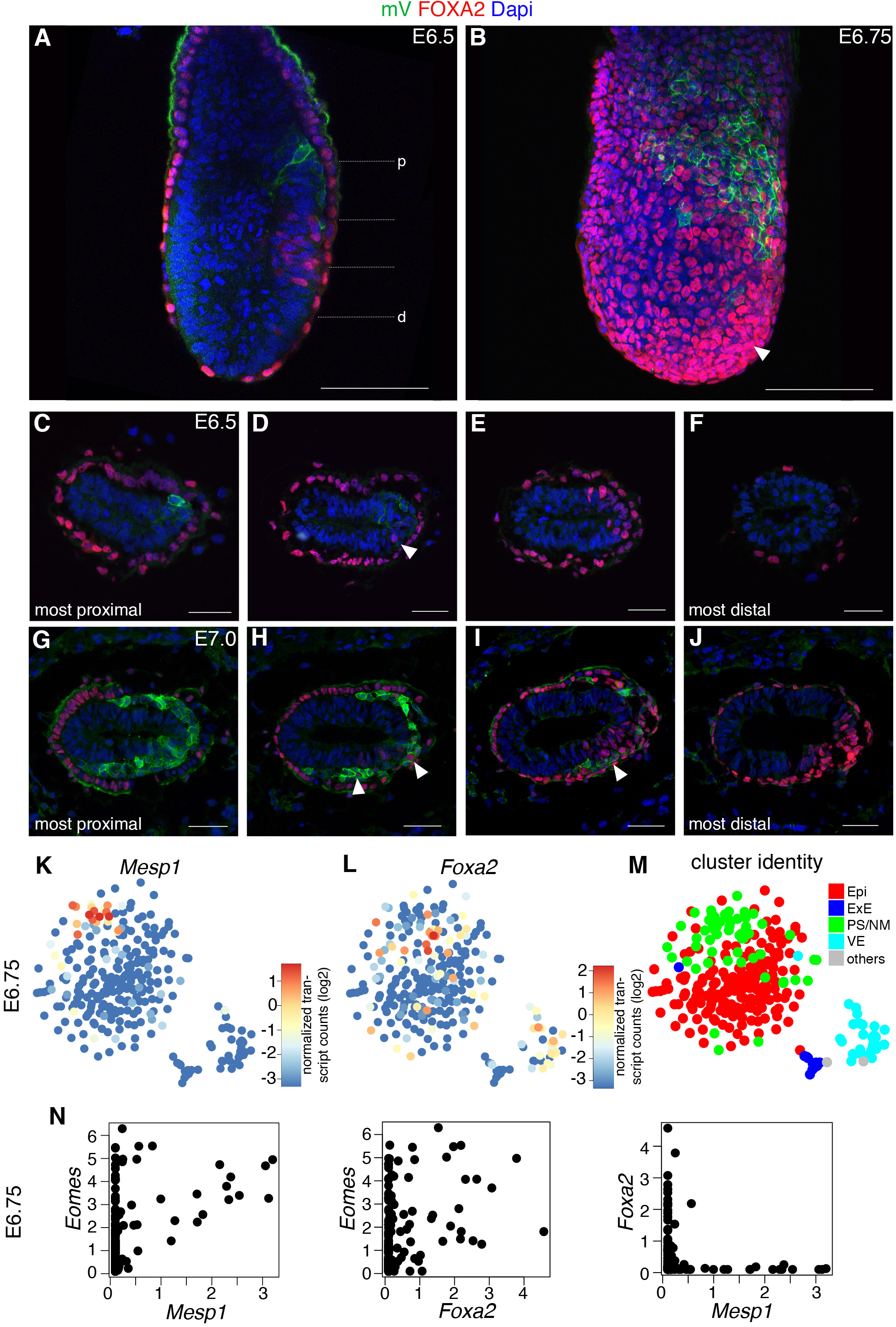
Spatial separation of *Eomes* dependent *Mesp1*^*mV*^ labelled AM and FOXA2 positive DE progenitors in the posterior epiblast and PS. **A-J**) Immunofluorescence staining in *Mesp1*^*mV*^ embryos using anti-GFP (green) and anti-FOXA2 (red) antibodies (n ≥ 3 embryos). FOXA2 is present in the cells of the VE. **A**) Sagittal section of an E6.5 *Mesp1*^*mV*^ embryo showing proximal *Mesp1*^*mV*^ positive cells and distal FOXA2 positive cells in the posterior epiblast (n = 1 embryo). **B**) Maximum intensity projection (MIP) of an E6.75 *Mesp1*^*mV*^ embryo (n = 2 embryos). *Mesp1*^*mV*^ positive cells are present in the proximal region of the embryo. FOXA2 positive cells in the VE are covering the whole embryo. In the posterior distal embryo FOXA2 positive cells are especially dense, probably corresponding to newly generated FOXA2 positive DE progenitors (arrowhead). **C-J**) Transverse sections at different levels of E6.5 (**C-F**) and E7.0 (**G-J**) *Mesp1*^*mV*^ embryos. **C** and **G** show the most proximal and **F** and **J** show the most distal sections. The approximate levels of the transverse sections are indicated in **A**, along the proximal (p) to distal (d) axis. Proximal sections contain *Mesp1*^*mV*^ positive cells and the more distal sections contain FOXA2 positive cells. A single FOXA2 positive cell in the *Mesp1*^*mV*^ positive domain of the epiblast in (**D**) is indicated with an arrowhead. At E7.0 there is an intermediate zone of mixed *Mesp1*^*mV*^ and FOXA2 positive cells (**H** and **I**). Few *Mesp1*^*mV*^/FOXA2 double positive cells are present (**H** and **I** arrowheads). Scale bars represent 100 μm (sagittal section and MIP) and 50 μm (transverse sections). **K, L**) t-SNE representation of E6.75 scRNA-seq data showing the expression of *Mesp1* (**K**) and *Foxa2* (**L**). The scale represents log2 normalized transcript counts. M) t-SNE plot with assigned identities to different clusters at E6.75. Anterior mesoderm (AM), epiblast (Epi), extraembryonic ectoderm (ExE), primitive streak (PS), visceral endoderm (VE). **N**) Scatter plots of single cells at E6.75 indicating *Eomes/Mesp1*, *Eomes/Foxa2* and *Foxa2/Mesp1* expression. Only one single *Foxa2/Mesp1* double positive cell is detected. X- and y-axes indicate normalized transcript counts.

The simultaneous analysis of *Mesp1*^*mV*^ and FOXA2 showed that AM and DE progenitors are generated at distinct domains along the PS (Fig. 3A-J). At E6.5 FOXA2 positive DE progenitors in the posterior epiblast are located more distally and still remain within the epithelial epiblast in comparison to proximally located *Mesp1*^*mV*^ positive AM cells, of which some have already delaminated from the PS as observed in sagittal (Fig. 3A), or in consecutive transverse sections (Fig. 3C-F). Additionally, FOXA2 broadly marks the cells of the VE (Fig. 3A-J). The distribution of proximally located AM and distal DE progenitors was also found at E6.75 (Fig. 3B) and E7.0 (Fig. 3G-J), where the most proximal sections showed *Mesp1*^*mV*^ expression in the PS and in cells of the mesoderm layer (Fig. 3G). More distal regions of the PS contained a mix of *Mesp1*^*mV*^ and FOXA2 single positive cells. The vast majority of cells in the mesoderm layer were *Mesp1*^*mV*^ expressing, and only rarely *Mesp1*^*mV*^ positive cells that also showed a FOXA2 signal were found (Fig 3H,I, arrowheads). These cells are located at the mid-level along the proximo-distal axis where the domains of *Mesp1*^*mV*^ positive cells and of FOXA2 positive cells meet and they most likely represent recently described FOXA2 positive progenitors that contribute to the cardiac ventricles (Bardot et al. 2017). Their location only at the border of *Mesp1* and *Foxa2* positive areas suggests that these are specialized cells rather than bipotential progenitors. At the distal tip of the PS only FOXA2 positive cells were found within the streak and all cells that have left the PS were FOXA2 positive (Fig. 3J). In summary, AM and DE progenitors are generated in mostly non-overlapping domains and cells are already lineage-separated when they are still located within the epithelial epiblast at the level of the PS.

Next, we employed scRNA-seq to analyze the segregation of *Foxa2* expressing DE and *Mesp1* expressing AM progenitors by their RNA expression profiles (Fig. 3K-N). At E6.75 and E7.5 *Mesp1* and *Foxa2* expression is found in distinct cells on the t-SNE maps (Fig. 3K-M and Fig. S2A-C). As expected, plotting cells for their expression of *Eomes*, *Mesp1* and *Foxa2* at E6.75 shows co-expression of *Mesp1* or *Foxa2* with *Eomes* (Fig. 3N, first and second plot). Notably, the *Mesp1* and *Foxa2* expressing cells are mutually exclusive, with the exception of one observed *Mesp1/Foxa2* double positive cell (Fig. 3N, third plot), which is also in accordance with the immunofluorescence staining analysis (Fig. 3A-J). At E7.5 *Eomes* starts to be rapidly downregulated and consequently increasing numbers of *Mesp1* or *Foxa2* single positive, *Eomes* negative cells are found (Fig. S2D, Tables S3, S4), and *Mesp1* positive and *Foxa2* positive cell populations remain exclusive (Fig. S2D). To confirm our analysis we also employed a published scRNA-seq data set that contains larger cell numbers (Pijuan-Sala et al. 2019). Here, we included and combined timepoints from E6.5 to E7.5 (E6.5, E6.75, E7.0, E7.25, E7.5) and performed cell clustering using the Seurat package (Stuart et al. 2019). Very similar clusters were identified as from our dataset (Fig. S3A, B) and similarly to the analysis of our dataset, the *Mesp1* and *Foxa2* expression domains on the UMAP representations were largely non-overlapping (Fig. S3D, E). Also, plotting the cells for their expression values at separate timepoints shows that *Mesp1* positive and *Foxa2* positive cell populations are mostly exclusive (Fig. S4). Quantification of *Mesp1/Foxa2* double positive cells within both datasets shows that more than 95% of *Mesp1* or *Foxa2* positive cells are single positive and only between 1.7 to 5% of cells at different timepoints are double positive (Fig. S3F and Fig. S4) demonstrating the separation of the progenitor populations of AM and DE.

Interestingly, at E6.75 the *Mesp1* positive cells cluster closely together on the t-SNE map, whereas *Foxa2* positive cells are found more scattered within the clusters of Epi/PS/NM (Fig. 3K, L, M and Fig. S2E). This suggests that *Foxa2* positive cells have less homogenous expression profiles which are more similar to unspecified epiblast cells at these early timepoints of analysis, since at E7.5 *Foxa2* positive cells form discrete clusters of node and DE cells (Fig. S2B, C, and F).

In summary, the simultaneous analysis of early emerging AM and DE progenitors at E6.5 reveals the spatial separation of their sites of origin within the population of *Eomes* expressing cells. *Mesp1*^*mV*^ mesoderm progenitors are generated more proximally and leave the PS earlier than distally generated *Foxa2* positive DE progenitors. ScRNA-seq analysis shows that the progenitor populations of AM and DE are separated as shown by the exclusive expression of early markers, *Mesp1* and *Foxa2*.

### Eomes dependent AM progenitors are specified at earlier timepoints than DE progenitors

Our analyses and published literature show that the generation of mesoderm and DE progenitors is spatially separated along the forming PS (Fig. 3; (Lawson and Pedersen 1987; Tam and Beddington 1987; Lawson et al. 1991; Tam and Behringer 1997). The fact that the PS elongates over time in a proximal to distal fashion suggests that mesoderm and DE progenitor specification is also temporally separated. To test the temporal sequence of lineage specification downstream of *Eomes*, we performed time dependent genetic lineage tracing using a tamoxifen inducible *Eomes*^*CreER*^ mouse line expressing CreER from the *Eomes* locus (Pimeisl et al. 2013) in combination with a Cre inducible fluorescent reporter ((Muzumdar et al. 2007), Fig. 4A). This *Rosa26*^*mTmG*^ reporter strain ubiquitously expresses membrane bound Tomato that switches to membrane bound GFP following Cre recombination. Short-term administration of tamoxifen (90 min) to dissected and morphologically staged embryos in culture was used to label *Eomes* positive cells at different developmental timepoints from E6.25 – E7.5 (Fig. 4A). Embryos were sorted into three groups according to the stage at the time of dissection (E6.25-E6.5, E6.75-E7.0, E7.25-7.5), tamoxifen treated for 90 minutes and cultured for additional 24 hours. Whole embryos were imaged as z-stacks to evaluate if the presence of GFP labeled cells within the mesoderm and endoderm layers depends on the timepoint of Cre induction (Fig. 4A and B). Of note, in addition to labeling of epiblast derived cell types, this approach also marks VE cells, since *Eomes* expression is also found here (Fig. 4C).

**Figure 4:**
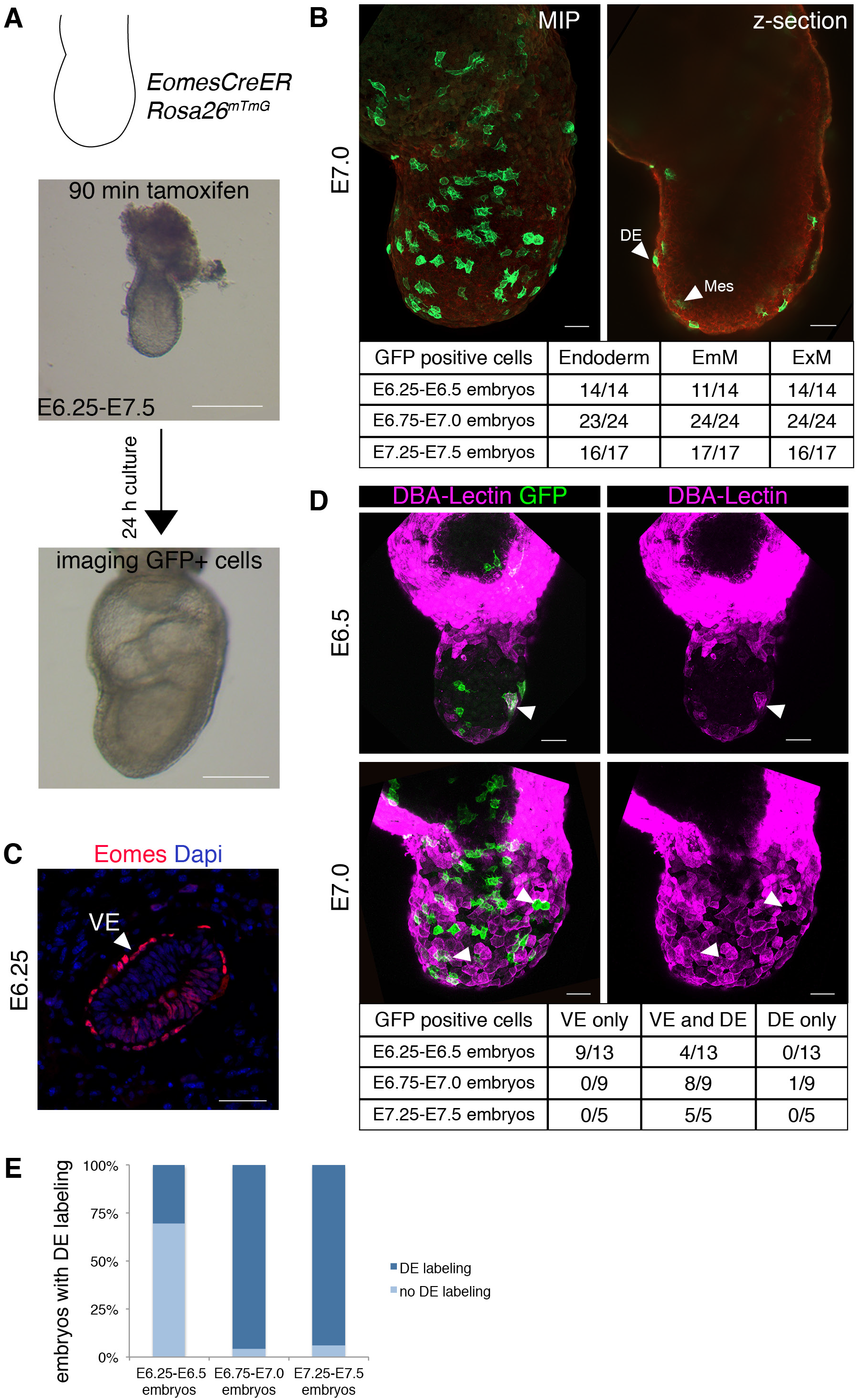
AM and DE are specified from *Eomes* positive cells following a sequential temporal order. **A**) Schematic of time dependent lineage tracing in E6.25 to E7.5 embryos carrying the *Eomes*^*CreER*^ and the *Rosa26*^*mTmG*^ reporter alleles (Muzumdar et al. 2007). Embryos were dissected, staged, and treated for 90 minutes with tamoxifen to induce CreER activity, followed by culture for 24 hours without tamoxifen and 3D imaging. A total number of 55 embryos were analyzed. **B**) Maximum intensity projection (MIP) and an optical section of an exemplary embryo treated with tamoxifen at E7.0. The MIP was used for the identification of all GFP positive cells, while the germ layer position of GFP positive cells was evaluated in optical sections. The contribution of *Eomes* expressing cells from labeling at different timepoints to different cell types is summarized in the table below the images. **C**) Transverse section of an E6.25 embryo stained with anti-EOMES antibody shows EOMES in the VE and in the epiblast. **D**) MIPs of embryos stained for DBA-lectin to identify VE cells. The upper panel shows an embryo treated with tamoxifen at E6.5. Lectin staining (arrowhead) of GFP positive cells indicates VE cells. The other GPF positive cells are in the mesoderm layer. The lower panel shows an embryo that was treated with tamoxifen on E7.0 and GFP positive cells in the endoderm layer are of both VE (lectin positive) and DE (lectin negative) origin as shown by arrowheads. The table summarizes the amounts of embryos with contribution of GFP positive cells to VE or DE, or both. 27 embryos were reanalyzed for DBA-lectin staining. **E**) Bar graph representing the percentage of embryos with GFP labeling in the DE. 70% of early E6.25 labeled embryos do not show GFP positive cell contribution to the DE lineage. All later timepoints of labeling there is a robust contribution of labeled cells to the DE lineage. Scale bars are 200 μm in **A**) and 50 μm in all other panels.

In a first analysis, 53 of 55 embryos showed labeling of both endoderm and mesoderm (including EmM and ExM) (Fig. 4B). Interestingly, three E6.25 embryos expressed GFP only within the ExM, supporting the observation that ExM is the first cell population generated in the PS (Parameswaran and Tam 1995; Kinder et al. 1999). As we were interested in the DE population within the labeled cells of the endoderm layer originating from *Eomes* positive cells in the posterior epiblast/PS we needed to discriminate VE from DE cells. Therefore we additionally stained all E6.25-E6.5 labeled embryos and some of the older embryos with the lectin dolichos biflorus agglutinin (DBA-lectin) that specifically labels VE cells but not DE (Fig. 4D) (Kimber 1986). This revealed that the GFP positive cells in the endoderm layer of 9 out of 13 (69%) E6.25-E6.5 labeled embryos were exclusively of VE origin, indicating that no DE is formed yet in most of the E6.25-E6.5 embryos (Fig. 4D). All E6.25-E6.5 embryos had GFP positive cells in mesoderm cells (EmM 10/13) (Fig. 4B). The GFP positive cells in the endoderm layer of the remaining 4 E6.25-E6.5 labeled embryos were of mixed DE and VE origin. Thus, we confirm the existence of a short time window before E6.5 during which *Eomes* expressing cells in the posterior epiblast give rise to mesoderm (Fig. 4D, E). Starting from E6.5 progenitors of mesoderm and DE are both present (Fig. 3) and therefore embryos that were tamoxifen treated at E6.5-E6.75 or later showed GFP labeling both in mesoderm and DE cells (Fig. 4D, E). In summary, lineage tracing of *Eomes* positive cells in E6.25-E6.5 embryos shows robust mesoderm labeling but absence of DE labeling in 70 % of the embryos, while at later stages all embryos show labeling of both mesoderm and DE cells (Fig. 4B, E). These experiments thus confirm that mesoderm and DE specification is also temporally separated so that mesoderm progenitor specification slightly proceeds DE formation.

### Eomes expressing epiblast cells differentiate directly to either AM or DE lineages

As the RaceID algorithm did not identify distinct progenitor populations for AM and DE within the *Eomes* positive, lineage marker negative cells of the epiblast (Fig. 1I, L), we wanted to investigate scRNA-seq expression profiles during this lineage segregation in more detail. Thus, we analyzed the single cell transcriptomes of *Eomes* positive cells (expression cut-off 0.3 normalized transcript counts) from the datasets with larger cell numbers (Pijuan-Sala et al. 2019) at timepoints from E6.5 to E7.5. According to our analysis (Fig. 1, 3) the *Eomes* positive population should include the unspecified progenitors as well as early AM and DE progenitors. Extraembryonic cells were excluded from the analysis. VarID (Grün 2020) identified cell clusters representing the posterior epiblast and two branches consisting of the proximal PS, NM, AM and ExM and of the distal PS, axial mesoderm (AxM), node, and DE (Fig. 5A and Fig. S5A).

**Figure 5:**
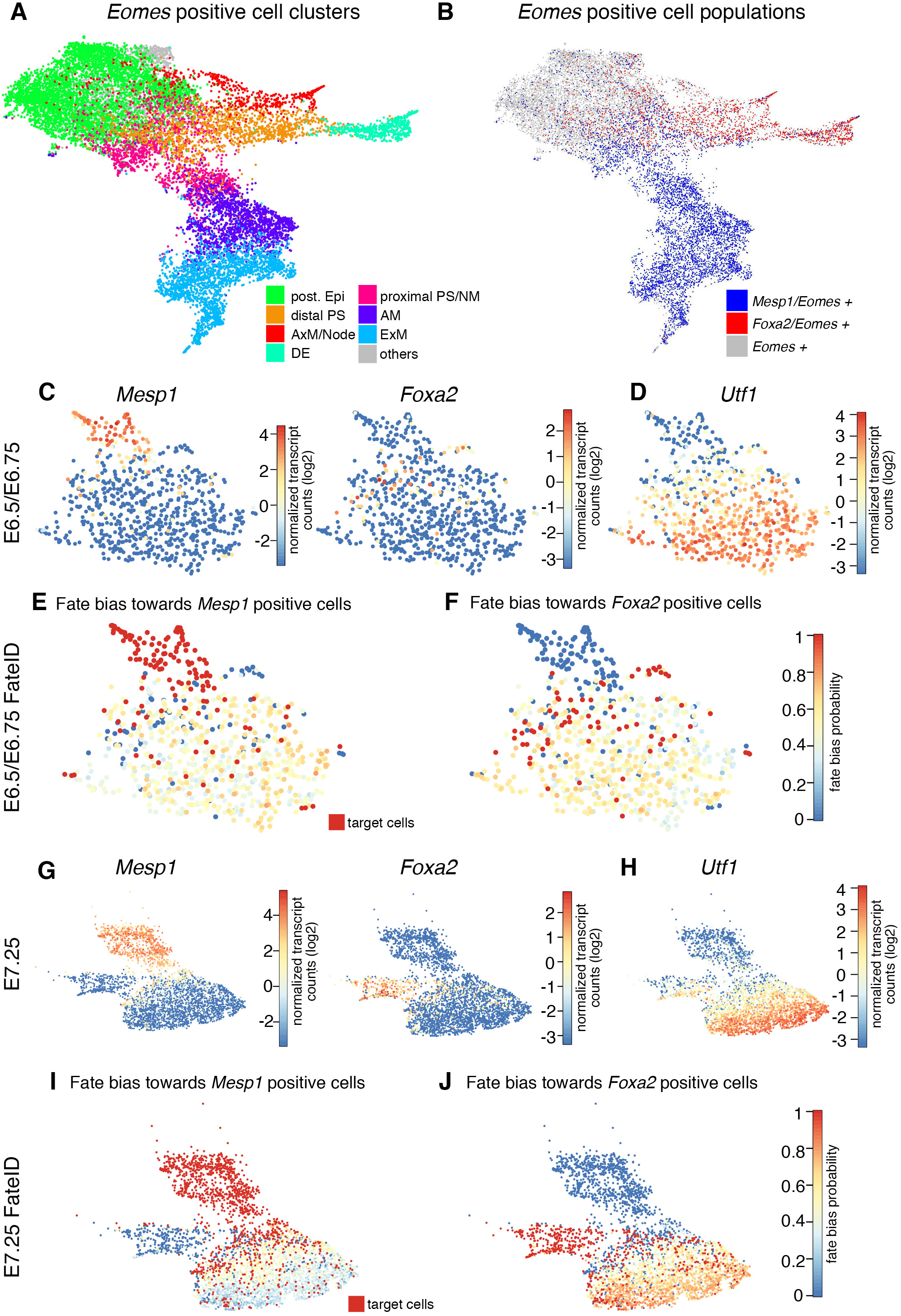
*Eomes* positive posterior epiblast cells directly differentiate to either AM or DE. **A**) UMAP representation of all *Eomes* positive cells from E6.5 to E7.5 embryos with an expression cut off >0.3 counts after the exclusion of extraembryonic cells (ExE and VE) (data from (Pijuan-Sala et al. 2019). Assigned clusters identities are indicated. Anterior mesoderm (AM), axial mesoderm (AxM), definitive endoderm (DE), posterior epiblast (post. Epi), nascent mesoderm (NM), extraembryonic mesoderm (ExM), primitive streak (PS). **B**) UMAP representation of *Mesp1/Eomes* double positive cells (blue), *Foxa2/Eomes* double positive cells (red), and *Eomes* single positive cells (grey) within all *Eomes* positive cells. The cut off for *Mesp1* and *Foxa2* expression was set to >0.3 counts. **C**, **D**) *Eomes* positive cells from E6.5/E6.75 were clustered and *Mesp1* and *Foxa2* (**C**) and *Utf1* (**D**) expression was plotted onto the UMAP representation. **E**, **F**) FateID analysis of embryonic *Eomes* positive cells from timepoints E6.5 and E6.75. The fate bias probability is indicated in single *Eomes* positive cells towards *Mesp1* positive cells (**E**) and *Foxa2* positive cells (**F**) (red, target cells). **G**, **H**) *Eomes* positive cells from E7.25 were clustered and *Mesp1* and *Foxa2* (**G**), and *Utf1* (**H**) expression was plotted onto the UMAP representation. **I, J**) FateID analysis of embryonic *Eomes* positive cells at timepoint E7.25. The fate bias probability is indicated in single *Eomes* positive cells towards *Mesp1* positive cells (**I**) and *Foxa2* positive cells (**J**) (red, target cells). Color scale represents fate bias probabilities on the scale from 0 to 1. Scale bar for gene expression on UMAP maps represents log2 normalized transcript counts.

These embryonic *Eomes* positive cells were categorized into three groups of *Mesp1/Eomes* double positive cells (blue), *Foxa2/Eomes* double positive cells (red), and *Eomes* single positive cells (grey) (expression cut-off 0.3 normalized transcript counts for all three genes) (Fig. 5B). Differential gene expression analysis between the *Eomes*/*Mesp1* or *Eomes*/*Foxa2* double positive cells and *Eomes* single positive cells showed that *Foxa2* positive cells expressed higher endoderm and axial mesoderm marker genes (e.g. *Sox17*, *Cer1*, *Gsc*) and *Mesp1* positive cells showed increased expression of mesodermal/mesenchymal/EMT genes (e.g. *Fn1*, *Lefty2*, *Myl7*, *Snai1*) (Fig. S5B and Table S5). Both *Mesp1* and *Foxa2* positive cells showed a downregulation of epiblast markers (e.g. *Pou3f1*, *Utf1*, *Slc7a3*) (Fig. S5B and Tables S5). This indicates that these two cell populations start to differentiate towards their respective fates. Overall more genes were differentially regulated in *Mesp1* positive cells than in *Foxa2* positive cells (126 vs. 43 genes >2-fold changed), indicating that DE progenitors are more similar in their expression profile to cells of the epithelial epiblast or that AM progenitors are further differentiated.

To investigate if there exists a differentiation bias towards either AM or DE progenitors already within *Eomes* single positive cells we used FateID, an algorithm that quantifies the fate bias in progenitor cell populations with known trajectories (Herman et al. 2018). We analyzed each timepoint separately to avoid artefacts originating from different developmental stages of cells, with the exception of cells from E6.5 and E6.75 that were combined to increase cell numbers (Fig. 5C, G and Fig. S5D). To perform FateID lineage bias analysis we defined early *Eomes*/*Mesp1* and *Eomes*/*Foxa2* double positive cells as target cells and excluded more differentiated clusters (target cells, shown in red cells in Fig. 5E, F, I, J, and Fig. S5F, G). On the respective UMAP representations of E6.5/E6.75 cells, early *Mesp1* positive cells are grouped and *Foxa2* positive cells are more scattered (as described in Fig. 3K, L; Fig. 5C), while at later time points *Eomes/Mesp1* and *Eomes/Foxa2* double positive cells form two distinct branches (Fig. 5G and Fig. S5D). *Utf1* expression is shown to indicate the undifferentiated epiblast population within the *Eomes* positive cells (Fig. 5D, H and Fig. S5E) (Tosic et al. 2019; Galonska et al. 2014). We then calculated the fate bias probabilities (between 0 and 1) of *Eomes* single positive cells for each target group, i.e. *Eomes/Mesp1* positive cells or *Eomes/Foxa2* positive cells in red (fate bias probability of 1). The target cells of the other fate appear blue (fate bias probability of 0). This analysis showed that at E6.5/E6.75 *Eomes* single positive cells have a similar fate bias probability towards both lineages of AM and DE (yellow cells, Fig. 5E, F) and thus did not exhibit a clear fate-bias towards either lineage. Accordingly, only few differentially expressed genes were found when we compared expression values in mesoderm and endoderm biased cells (fate bias probability cut off was set to 0.6 for each of the respective lineages) and these were mostly not known early lineage marker genes (12 genes >2-fold changed, Fig. S5C and Table S6). At E7.25 the *Eomes* single positive cells of the more undifferentiated epiblast (Fig. 5H, *Utf1* positive) were biased towards *Eomes/Foxa2* positive target cells (orange cells, Fig. 5J). The cells closer to the branching point mostly did not display a clear fate bias showing an intermediate probability for both lineages (yellow cells, Fig. 5I, J). This indicates that at E.725 most mesoderm downstream of *Eomes* has already been generated and the majority of the remaining *Eomes* single positive epiblast cells will give rise to DE/axial mesoderm. Analysis of differentially expressed genes between endoderm and mesoderm biased cells at E7.25 reveals that epiblast markers are more strongly expressed in endoderm fated cells whereas mesoderm fated cells already show a pronounced mesodermal expression profile (52 genes >2-fold changed, Fig. S5C and Table S6). At E7.0 FateID analysis resulted in an intermediate result between E6.5/E6.75 and E7.25 (Fig. S5E), showing the progression of fate bias towards *Foxa2* positive population in the *Eomes* expressing epiblast cells over time (Fig. S5F, G).

In conclusion, until E7.0 the *Eomes* single positive posterior epiblast cells are not fate biased towards either lineage before the onset of *Mesp1* or *Foxa2* expression. *Mesp1* and *Foxa2* are expressed in distinct cell populations and there is no evidence of a cell population co-expressing lineage markers for mesoderm and DE. This suggests that cells differentiate directly from a posterior epiblast state to either mesoderm or endoderm lineages.

## Discussion

To date the understanding of lineage specification on the level of individual cells within the gastrulation stage embryo remains limited. It is still unclear how cells in close proximity acquire different fates according to local signaling environments and how these specification events are regulated in a temporal manner. In this study we have analyzed the emergence of AM and DE populations that are both dependent on activities of the transcription factor EOMES (Arnold et al. 2008; Costello et al. 2011; Teo et al. 2011). Interestingly, we observed that almost all cells passing through the PS during the first day of gastrulation (E6.5 to E7.5) are positive for EOMES. These cells give rise to the mesoderm derivatives of the anterior embryo and the entire DE as previously shown by lineage tracing of *Eomes* positive cells (Costello et al. 2011; Arnold et al. 2008). The *Eomes* positive population in the early PS/mesendoderm was thought to be one of several populations leaving the PS between E6.5 and E7.5 (Robertson 2014). However, our data indicate that early posterior epiblast cells uniformly express *Eomes*. These results thus indicate that between E6.5 and E7.5 only cells that will contribute to the mesoderm of the anterior embryo and the DE progenitors leave the PS and posterior mesodermal tissues are generated after E7.5. Accordingly, spatial gene regulatory network analysis of gastrulation stage embryos indicates that separate anterior and posterior mesoderm populations exist at E7.5 (Peng et al. 2019). Our FateID analysis further indicates that AM downstream of *Eomes* is mainly generated until E7.25. During following stages mesoderm formation is most likely regulated by other factors such as the related T-box factor *T/Brachyury* and WNT signaling (Koch et al. 2017; Wymeersch et al. 2016). The existence of distinct anterior and posterior mesoderm populations downstream of different T-box factors has been suggested previously and is also observed in the zebrafish (Kimelman and Griffin 2000). The molecular details of this transition in the regulation of gastrulation are currently incompletely understood.

The first lineage decision following *Eomes* expression in epiblast cells segregates AM and DE. Here, we show that the AM and DE marker genes, *Mesp1* and *Foxa2*, are already expressed in epithelial epiblast cells at the PS and are mostly exclusive. Therefore, we can place the event of lineage segregation within the PS before cells migrate to form the mesoderm layer. These results are in agreement with previous observations of a separation of T and FOXA2 protein expression domains in E6.5 embryos (Burtscher and Lickert 2009). Also, earlier cell tracing experiments have shown that cells are restricted in their potency after their passage through the PS (Tam et al. 1997). Live embryo imaging analysis has shown that DE and mesoderm progenitors leave the PS and migrate together within the mesodermal wings before DE cells insert into the outer VE layer (Viotti, Nowotschin, et al. 2014). As we show that DE cells are already specified as they leave the PS, it will be interesting to investigate how DE cells behave within the mesoderm population and which mechanisms are used to separate them during the anterior-ward migration.

During early gastrulation AM and ExM subtypes as well as most of the DE cells are generated (Parameswaran and Tam 1995; Lawson et al. 1991; Tam et al. 2007; Lawson et al. 1986; Lawson and Pedersen 1987; Kinder et al. 1999). Here, we show that mesoderm and DE are produced at distinct places along the PS and that mesoderm is specified earlier than DE as summarized in a model (Fig. 6). The proximal domain of the PS generates only mesoderm from the initiation of gastrulation. With a slight temporal delay the most distal tip of the PS produces only *Foxa2* positive DE and axial mesoderm progenitors, however, in this study we did not address the generation and distribution of the axial mesoderm. The intermediate PS regions generate both mesoderm and endoderm progenitors and here we also find some rare *Mesp1/Foxa2* double positive cells. This data fits well with earlier cell labeling experiments that had demonstrated that the most proximal PS produces mostly mesoderm (Lawson et al. 1991; Parameswaran and Tam 1995; Kinder et al. 1999). At E6.5 mesodermal progenitors already leave the PS, while DE progenitors are present in the more distal epiblast. To date, the evidence for delayed specification of the DE was derived from experiments showing that DE progenitor cells appear in the outer VE endoderm layer only at about E7.0 (Lawson and Pedersen 1987; Lawson et al. 1991). Using genetic timed lineage tracing we now demonstrate the time delay of DE cell specification in comparison to mesoderm, so that first DE cells are specified about 0.25 days after the mesodermal cells at E6.5 as also inferred from transcriptional analysis (Peng et al. 2019; Pijuan-Sala et al. 2019).

**Figure 6:**
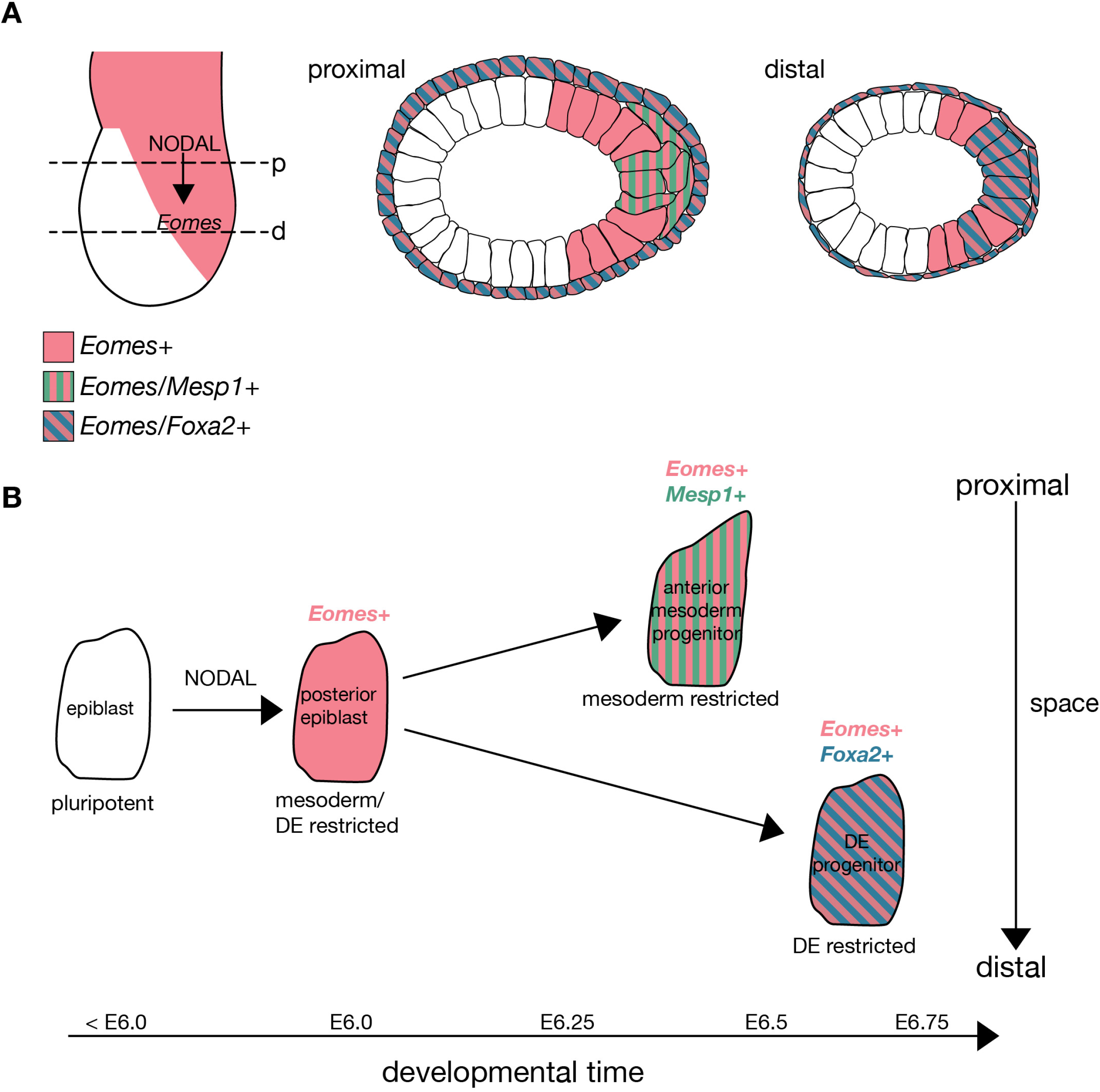
Model of spatial and temporal separation of DE and AM lineage specification downstream of *Eomes*. **A**) Induction of *Eomes* by NODAL/SMAD2/3 signals leads to the specification of AM and DE lineages. AM marked by *Mesp1* expression is generated in the proximal PS and *Foxa2* positive cells give rise to DE in the distal PS. The schematic shows an E6.5 embryo where few AM progenitor cells have already delaminated from the PS, while DE progenitor cells are still entirely located in the epiblast. The VE is positive for *Foxa2*. Proximal (p) and distal (d). **B**) Both AM and DE progenitor cells are specified from an unbiased *Eomes* positive progenitor cell at different localizations along the proximo-distal axis and at different timepoints. *Mesp1* and *Foxa2* indicate the first fully specified AM and DE cells, respectively.

Analysis of the fate bias of *Eomes* positive cells, which are not yet expressing *Mesp1* or *Foxa2* markers, indicates that an unbiased posterior epiblast state directly progresses to either AM or DE. In our scRNA-seq analysis we did not find evidence for progenitors co-expressing markers of both lineages, arguing for fast acting control mechanisms that independently promote AM or DE programs (Fig. 6B). However, we can’t rule out the existence of different, already lineage restricted progenitors for AM and DE, since we demonstrate that these are spatially separated cell populations. Embryonic clonal lineage analyses had suggested that the common posterior epiblast progenitor for mesoderm and DE represents only a very transient cell population since clones containing both AM and DE cells were only very rarely detected by genetic or labeled lineage tracing (Tzouanacou et al. 2009; Lawson et al. 1991). Novel approaches using a combination of scRNA-seq and molecular recording of cell lineage might be able to provide information on the lineage segregation and relationship of AM and DE in more detail, which was not explored in existing data sets to date (M. M. Chan et al. 2019).

Genetic experiments in the mouse embryo and cell differentiation studies in culture have demonstrated that NODAL/SMAD2/3 signals are major regulators in the AM and DE lineage decision (Dunn et al. 2004; Vincent et al. 2003; van den Ameele et al. 2012; Costello et al. 2011). Accordingly, elevated levels of NODAL signaling are required for DE specification, while AM is already formed at lower NODAL signaling levels. How the signaling thresholds of NODAL/SMAD2/3 signals are interpreted during this lineage choice remains unresolved. Our study supports previous observations (van Boxtel et al. 2015; Sako et al. 2016) that NODAL/SMAD2/3 signal interpretation might involve a temporal signal integration leading to delayed formation of DE in comparison to AM. Alternatively, additional signals might contribute to this lineage choice, such as WNT or FGF signals as previously suggested in zebrafish (van Boxtel et al. 2018) and ES cells (Wang et al. 2017).

In conclusion, this study demonstrates that the generation of the *Eomes* dependent lineages of AM and DE is spatiotemporally separated during early gastrulation. These cells are molecularly separated early during the differentiation process and share as last common progenitor the *Eomes* expressing posterior epiblast cells.

## Materials and Methods

### Generation of the *Mesp1*^*mVenus*^ allele

To generate a fluorescent allele to follow *Mesp1* expressing cells during gastrulation we targeted the *Mesp1* locus by homologous recombination to insert a membrane bound Venus fluorescent protein reporter (mVenus) into the locus. Following the mVenus coding sequence the *Mesp1* coding sequence including a 3xFLAG C-terminal tag was inserted. These two coding sequences are linked by a T2A peptide, which leads to co-translational cleavage of the two proteins, resulting in independent mVenus and Mesp1-3xFLAG proteins (Fig. 2A). The start site of mVenusT2AMesp1-3xFLAG was inserted at the translational start site of the *Mesp1* gene. The 5’ homologous arm spans from an AfeI site upstream of *Mesp1* exon 1 to the *Mesp1* translational start site. The mVenusT2AMesp1-3xFLAG sequence followed by a bGH PolyA signal was then inserted via the 5’UTR EcoRI site and a FspA1 site within *Mesp1* exon1, thereby deleting 337 bp of the *Mesp1* CDS. A PGK-neomycine resistance (neoR) cassette flanked by loxP sites was inserted downstream of mVenusT2AMesp1-3xFLAG. Between the mVenusT2AMesp1-3xFLAG insert and the neoR cassette a NdeI site was introduced for screening by southern analysis. The 3’ homology region spans to a NsiI site downstream of *Mesp1* exon2 and was flanked by a pMCI-TK negative selection cassette.

Linearized targeting vector was electroporated into CCE ES cells, and neomycine and FIAU resistant ES cell clones were screened by genomic southern blot. Genomic DNA was digested with NdeI and probed with an external 3’ probe (wild type (wt) allele: 8.4 kb, mutant allele: 4.7 kb, Fig. 2B). Two independent positive clones were injected at morula stage for chimera generation. *Mesp1*^*mVenus*^ mice were genotyped by PCR at 62°C annealing temperature to detect the wild type allele (334 bp) and the knock-in allele (419 bp) using the following primers: wt forward primer 5’-CGCTTCACACCTAGGGCTCA -3’; wt reverse primer 5’-TGTGCGCATACGTAGCTTCTCC -3’; ki forward primer 5’-GCCAATGCAATCCCGAAGTCTC-3’; ki reverse primer 5’-GCCCTTGGACACCATTGTCTTG -3’ (Fig. 2C). The neomycine cassette was removed by crossing *Mesp1*^*Venus*^ positive males to females carrying the *Sox2::Cre* transgene (Vincent and Robertson 2003).

### Mice

*Mesp1*^*Venus*^ mice were backcrossed to the NMRI strain and otherwise kept as homozygotes, since they were viable and fertile and showed no obvious phenotypic differences to wt. *Eomes*^*mTnG*^ mice were also kept on a NMRI background and were kept as heterozygotes. Mice were maintained as approved by the Regierungspräsidium Freiburg (license numbers G11/31 and X19/O2F).

### Whole mount immunofluorescence of embryos

Embryos were dissected in PBT (PBS with 0,1% Tween-20), fixed for one hour in 4% PFA in PBS at 4°C or on ice and washed twice in PBT. At this point embryos can be kept at 4°C for at least a month. To perform the staining embryos were permeabilized in 0.3% Triton X-100 in PBT at RT for 30 min. Embryos were blocked for 2 hours at RT in blocking solution (1% BSA in PBT). The primary antibodies (for list of antibodies see ‘immunofluorescence on embryo sections’) were incubated in blocking solution at 4°C over night. Embryos were washed 4 × 5min in PBT at RT and then incubated with the secondary antibodies in blocking solutions for 3 hours at RT. Embryos were washed 2 × 5 min in PBT at RT and then stained with DAPI for 30 min at RT to overnight at 4°C and then washed with PBT. Embryos were stored and imaged in PBT. Imaging was performed on a Zeiss inverted laser-scanning microscope or a Zeiss spinning disk inverted microscope in glass bottom dishes.

### Immunofluorescence on embryo sections

Embryos were fixed in 4% PFA for one hour at 4°C in the deciduae that were opened to expose the embryo. Deciduae were washed with PBT and then processed through 15% and 30% sucrose/PBS at 4°C and incubated for at least one hour in embedding medium (15% sucrose/7.5% gelatin in PBS) at RT prior to embedding. 6-7 μm thick sections were cut with a Leica cryotome. To perform the immunofluorescence staining sections were washed 3 × 5 min in PBS and permeabilized in PBT containing 0.2% Triton-X100. Sections were blocked in blocking solution (1% BSA in PBT) for one hour at RT. The primary antibodies were added in blocking solution at 4°C over night. The slides were washed 3 × 5 min with PBS and then incubated with the secondary antibody in blocking solution for one hour at RT. After washing the antibody away with PBS, the sections were stained with DAPI in PBT for 5 min and then mounted with ProLong Diamond Antifade Mountant (Life technologies P36970) and imaged on an inverted Zeiss Axio Observer Z1 microscope. Primary antibodies used: GFP (1:1000, Abcam ab13970), RFP (1:500, Rockland 600-401-379), EOMES (1:300, Abcam ab23345), FOXA2 (1:500, Cell Signaling 8186). Secondary Alexa-Fluor-conjugated antibodies (Life Technologies) were used at a dilution of 1:1000.

### Time dependent lineage tracing using the *Eomes*^*CreER*^ allele

Embryos were isolated on E6 or E7 in prewarmed dissection medium (10% fetal calf serum (FCS) in DMEM/F12 containing Glutamax) and were then placed in embryo culture medium (50% DMEM/F-12 containing Glutamax and 50% Rat Serum) containing 10 μm of 4-OH-tamoxifen (Sigma H7904, dissolved at 10mM in DMSO) for 90 min. Embryos were washed 3x in dissection medium and placed individually in ibidi 8-well slides in embryo culture medium without 4-OH-Tamoxifen. Embryos were cultured for 24 hours in regular tissue culture incubators at 37°C with 5% CO_2_. A picture was taken before and after these 24 hours of each individual embryo. After the incubation embryos were fixed and stained with GFP and RFP antibodies as described above and imaged on a Zeiss spinning disk inverted microscope in glass bottom dishes.

### Fluorescent DBA-Lectin staining on whole mount embryos

Embryos from the lineage tracing experiments were re-stained with the biotinylated lectin dolichos biflorus agglutinin (DBA) (Sigma L6533). Because embryos were already stained with GFP and RPF antibodies, no extra blocking step was performed. Embryos were washed in PBT and then the DBA-lectin was added at a dilution of 1:1000 in PBS with 1% BSA at 4°C over night. The next day embryos were washed 3 × 10 min with PBT and then incubated with Alexa-Fluor-647-streptavidin (Molecular Probes, S21374, dissolved in PBS at 1mg/ml) in PBS containing 1% BSA at a dilution of 1:500 for one hour at RT. Before adding the streptavidin the tube was quickly centrifuged. Finally, embryos were washed 3x in PBT and imaged on a Zeiss inverted laser-scanning microscope in glass bottom dishes.

### Collection of embryo cells for single-cell RNA sequencing

Embryos were dissected in pre-warmed dissection medium (10% fetal calf serum (FCS) in DMEM/F12 containing Glutamax) and washed in pre-warmed PBS.

For the E6.75 timepoint the extraembryonic part was cut off and a picture of each embryo was taken and single embryos were transferred into the wells of a pre-warmed non-adhesive 96-well plate containing 40 μl of TrypLE Express (Gibco 12604013). The wells were coated with FCS before adding the TrypLE. Embryos were incubated at 37°C for 10 minutes with pipetting up and down once in between and at the end to make a single cell solution. Dissociation was stopped with 120 μl of dissection medium and cells were centrifuged for 2 min at 1000 rpm in the 96-well plate. The supernatant was removed and cells from one embryo were resuspended in 200 μl cold PBS. For hand-picking, the drop containing the cells was placed in a plastic petri dish. Cells were picked under a Leica M165 FC binocular using ES-blastocyst injection pipettes (BioMedical Instruments, blunt, bent ID 15 μm, BA=35°) and placed into 1.2 μl lysis buffer containing polyT primer with unique cell barcode. Embryos from the E7.5 timepoint were cut under to chorion to include the extraembryonic mesoderm in the analysis. The embryos were imaged and the embryos of one or two litters were pooled and processed in an FCS coated eppendorf tube in the same way as the E6.75 embryos. After centrifugation the cells were resuspended in 200 μl PBS and kept on ice until flow sorting.

### Single-cell RNA amplification and library preparation

Single-cell RNA sequencing of 576 hand-picked cells (E6.75) was performed using the CEL-Seq2 protocol while of 1152 flow-sorted cells (E7.5) was performed using the mCEL-Seq2 protocol (Hashimshony et al. 2016; Herman et al. 2018). Eighteen libraries with 96 cells each were sequenced per lane on Illumina HiSeq 2500 or 3000 sequencing system (pair-end multiplexing run) at a depth of ~200,000-250,000 reads per cell.

### Quantification of transcript abundance

Paired end reads were aligned to the transcriptome using bwa (version 0.6.2-r126) with default parameters (Li and Durbin 2010). The transcriptome contained all gene models based on the mouse ENCODE VM9 release downloaded from the UCSC genome browser comprising 57,207 isoforms, with 57,114 isoforms mapping to fully annotated chromosomes (1 to 19, X, Y, M). All isoforms of the same gene were merged to a single gene locus. Furthermore, gene loci overlapping by >75% were merged to larger gene groups. This procedure resulted in 34,111 gene groups. The right mate of each read pair was mapped to the ensemble of all gene loci and to the set of 92 ERCC spike-ins in sense direction (Baker et al. 2005). Reads mapping to multiple loci were discarded. The left read contains the barcode information: the first six bases corresponded to the unique molecular identifier (UMI) followed by six bases representing the cell specific barcode. The remainder of the left read contains a polyT stretch. For each cell barcode, the number of UMIs per transcript was counted and aggregated across all transcripts derived from the same gene locus. Based on binomial statistics, the number of observed UMIs was converted into transcript counts (Grün et al. 2014).

### Clustering and visualization of mCEL-Seq2 data

Clustering analysis and visualization of the data generated in this study were performed using the RaceID3 algorithm (Herman et al. 2018). The numbers of genes quantified were 19,574 and 20,108 in the E6.75 and E7.5 datasets, respectively. Cells with a total number of transcripts <3,000 were discarded and count data of the remaining cells were normalized by downscaling. Cells expressing >2% of *Kcnq1ot1*, a potential marker for low-quality cells (Grün et al. 2016), were not considered for analysis. Additionally, transcript correlating to *Kcnq1ot1* with a Pearson’s correlation coefficient >0.65 were removed. The following parameters were used for RaceID3 analysis: mintotal=3000, minexpr=5, outminc=5, probthr=10^−4^. Mitochondrial, ribosomal as well as genes starting with ‘Gm’ were excluded from the analysis. We observed strong batch effects in E6.75 dataset based on the day of the hand-picking. Batch effects were corrected by matching mutual nearest neighbors (MNNs) as described previously (Haghverdi et al. 2018). mnnCorrect function from the scran package was used for the batch correction (Lun et al. 2016). MNN-based batch correction was also performed on the combined *Eomes* positive dataset used for FateID analysis.

### Clustering and visualization of mouse gastrulation atlas data

Processed atlas data on mouse organogenesis from (Pijuan-Sala et al. 2019) was downloaded from ArrayExpress (accession number: E-MTAB-6967). The following time points and sequencing batches were analyzed: E6.5 (sequencing batch 1), E6.75 (sequencing batch 1), E7 (sequencing batches 1, 2 and 3), E7.25 (sequencing batch 2) and E7.5 (sequencing batches 1 and 2). Cells defined as doublets in the study were removed from the analysis. Integration of datasets from different time points and sequencing batches was performed using Seurat version 3 with default settings (Stuart et al. 2019). Ribosomal genes (small and large subunits) as well as genes with *Gm*-identifiers were excluded from the data prior to integration. The integrated dataset contained 45,196 cells. A focused analysis of *Eomes*-positive cells was performed using VarID (Grün 2020). From the complete dataset containing 45,196, cells with a total number of transcripts <6,000 were discarded and count data of the remaining cells were normalized by downscaling. Cells having normalized *Eomes* transcript counts >0.3 were considered as *Eomes*-positive (14,329 cells) and further clustered and visualized using VarID with the following parameters: large=TRUE, pcaComp=100, regNB=TRUE, batch=batch, knn=50, alpha=10, no_cores=20. Each batch contained cells from different time points and sequencing libraries. Dimensionality reduction of the datasets was performed using Uniform Manifold Approximation and Projection (UMAP).

### FateID analysis

In order to investigate the transcriptional priming of single *Eomes* positive cells towards the mesodermal and definitive endodermal (DE) fates, FateID (Herman et al. 2018) was run on the combined E6.75 and E7.5 dataset with cells having normalized *Eomes* transcript counts >0.3 using default parameters. *Mesp1* positive (mesoderm specified, normalized transcript count >0.3) and *Foxa2* positive (DE specified, normalized transcript count >0.3) cells were used as target cells. Extra-embryonic cells were excluded from the FateID analysis and t-distributed stochastic neighbor embedding was used for dimensional reduction and visualization of the results. Differential gene expression analysis was performed between cells biased towards one of the lineages with fate bias probability >0.5 using diffexpnb function. FateID analysis was also performed on the mouse gastrulation data (Pijuan-Sala et al. 2019) using the same parameters describe above but separately at the following different time points: E6.5/E6.75, E7.0, and E7.25. Uniform Manifold Approximation and Projection coordinates from the VarID analysis were used for the visualization of the results.

### Differential gene expression analysis

Differential gene expression analysis was performed using the diffexpnb function of RaceID3 algorithm. Differentially expressed genes between two subgroups of cells were identified similar to a previously published method (Anders and Huber 2010). First, negative binomial distributions reflecting the gene expression variability within each subgroup were inferred based on the background model for the expected transcript count variability computed by RaceID3. Using these distributions, a p value for the observed difference in transcript counts between the two subgroups was calculated and multiple testing corrected by the Benjamini-Hochberg method.

## Acknowledgments

We thank T. Bass for technical assistance and the staff of the Life Imaging Center of the Albert-Ludwigs University Freiburg for help with microscopy. We thank Shankar Srinivas for teaching Simone Probst mouse imaging techniques. We thank Matthias Weiß and the CEMT team for support with animal care and Katrin Schüle for critical reading of the manuscript.

## Competing interests

The authors declare no competing interests

## Funding

This study was supported by the German Research Foundation (DFG) through the Emmy Noether- and Heisenberg-Programs (AR 732/1-1/2/3, AR 732/3-1), project grant (AR 732/2-1,) project B07 of SFB 1140 (project ID 246781735), and project A03 of SFB 850 (project ID 89986987) to S.J.A, and by the Germany’s Excellence Strategy (CIBSS – EXC-2189 – Project ID 390939984) to D.G. and S.J.A. DG was supported by the Max Planck Society, the German Research Foundation (DFG) (SPP1937 GR4980/1-1, GR4980/3-1, and GRK2344 MeInBio), by the DFG under Germany’s Excellence Strategy (CIBSS – EXC-2189 – Project ID 390939984), by the ERC (818846 — ImmuNiche — ERC-2018-COG), and by the Behrens-Weise-Foundation.

## Data availability

The primary read files as well as expression count files for the single-cell RNA-sequencing datasets generated in this study are available to download from GEO (accession number: GSE151824).

## Notes

### Competing Interest Statement

The authors have declared no competing interest.

### Summary of Updates

Supplemental information was added

## References

Anders, S. and Huber, W., 2010. Differential expression analysis for sequence count data. Genome biology, 11(10), p.R106.

Ang, S.L. and Rossant, J., 1994. HNF-3 beta is essential for node and notochord formation in mouse development. Cell, 78(4), pp.561–574.

Ang, S.L. et al., 1993. The formation and maintenance of the definitive endoderm lineage in the mouse: involvement of HNF3/forkhead proteins. Development, 119(4), pp.1301–1315.

Arkell, R.M., Fossat, N. and Tam, P.P.L., 2013. Wnt signalling in mouse gastrulation and anterior development: new players in the pathway and signal output. Current Opinion in Genetics and Development, 23(4), pp.454–460.

Arnold, S.J. and Robertson, E.J., 2009. Making a commitment: cell lineage allocation and axis patterning in the early mouse embryo. Nature Reviews Molecular Cell Biology, 10(2), pp.91–103.

Arnold, S.J. et al., 2008. Pivotal roles for eomesodermin during axis formation, epithelium-to-mesenchyme transition and endoderm specification in the mouse. Development, 135(3), pp.501–511.

Baker, S.C. et al., 2005. The External RNA Controls Consortium: a progress report. Nature Methods, 2(10), pp.731–734.

Bardot, E. et al., 2017. Foxa2 identifies a cardiac progenitor population with ventricular differentiation potential. Nature Communications, 8, p.14428.

Brennan, J. et al., 2001. Nodal signalling in the epiblast patterns the early mouse embryo. Nature, 411(6840), pp.965–969.

Burtscher, I. and Lickert, H., 2009. Foxa2 regulates polarity and epithelialization in the endoderm germ layer of the mouse embryo. Development, 136(6), pp.1029–1038.

Chan, M.M. et al., 2019. Molecular recording of mammalian embryogenesis. Nature, 100(6298), p.64.

Chan, S.S.-K. et al., 2013. Mesp1 patterns mesoderm into cardiac, hematopoietic, or skeletal myogenic progenitors in a context-dependent manner. Cell Stem Cell, 12(5), pp.587–601.

Ciruna, B.G. and Rossant, J., 1999. Expression of the T-box gene Eomesodermin during early mouse development. Mechanisms of Development, 81(1–2), pp.199–203.

Conlon, F.L. et al., 1994. A primary requirement for nodal in the formation and maintenance of the primitive streak in the mouse. Development, 120(7), pp.1919–1928.

Costello, I. et al., 2011. The T-box transcription factor Eomesodermin acts upstream of Mesp1 to specify cardiac mesoderm during mouse gastrulation. Nature Cell Biology, 13(9), pp.1084–1091.

Dufort, D. et al., 1998. The transcription factor HNF3beta is required in visceral endoderm for normal primitive streak morphogenesis. Development, 125(16), pp.3015–3025.

Dunn, N.R. et al., 2004. Combinatorial activities of Smad2 and Smad3 regulate mesoderm formation and patterning in the mouse embryo. Development, 131(8), pp.1717–1728.

Frank, D.U. et al., 2007. System for inducible expression of cre-recombinase from the Foxa2 locus in endoderm, notochord, and floor plate. Developmental Dynamics, 236(4), pp.1085–1092.

Galonska, C., Smith, Z.D. and Meissner, A., 2014. In Vivo and in vitro dynamics of undifferentiated embryonic cell transcription factor 1. Stem cell reports, 2(3), pp.245–252.

Grün, D., 2020. Revealing dynamics of gene expression variability in cell state space. Nature Methods, 17(1), pp.45–49.

Grün, D. et al., 2016. De Novo Prediction of Stem Cell Identity using Single-Cell Transcriptome Data. Cell Stem Cell, 19(2), pp.266–277.

Grün, D. et al., 2015. Single-cell messenger RNA sequencing reveals rare intestinal cell types. Nature, 525(7568), pp.251–255.

Grün, D., Kester, L. and van Oudenaarden, A., 2014. Validation of noise models for single-cell transcriptomics. Nature Methods, 11(6), pp.637–640.

Haghverdi, L. et al., 2018. Batch effects in single-cell RNA-sequencing data are corrected by matching mutual nearest neighbors. Nature Biotechnology, 36(5), pp.421–427.

Hashimshony, T. et al., 2016. CEL-Seq2: sensitive highly-multiplexed single-cell RNA-Seq. Genome biology, 17(1), p.77.

Herman, J.S., Sagar and Grün, D., 2018. FateID infers cell fate bias in multipotent progenitors from single-cell RNA-seq data. Nature Methods, 15(5), pp.379–386.

Imuta, Y. et al., 2013. Generation of knock-in mice that express nuclear enhanced green fluorescent protein and tamoxifen-inducible Cre recombinase in the notochord from Foxa2 and T loci. genesis, 51(3), pp.210–218.

Kartikasari, A.E.R. et al., 2013. The histone demethylase Jmjd3 sequentially associates with the transcription factors Tb×3 and Eomes to drive endoderm differentiation. The EMBO journal, 32(10), pp.1393–1408.

Kimber, S.J., 1986. Distribution of lectin receptors in postimplantation mouse embryos at 6-8 days gestation. The American journal of anatomy, 177(2), pp.203–219.

Kimelman, D. and Griffin, K.J., 2000. Vertebrate mesendoderm induction and patterning. Current Opinion in Genetics and Development, 10(4), pp.350–356.

Kinder, S.J. et al., 1999. The orderly allocation of mesodermal cells to the extraembryonic structures and the anteroposterior axis during gastrulation of the mouse embryo. Development, 126(21), pp.4691–4701.

Kinder, S.J., Loebel, D.A. and Tam, P.P., 2001. Allocation and early differentiation of cardiovascular progenitors in the mouse embryo. Trends in cardiovascular medicine, 11(5), pp.177–184.

Kitajima, S. et al., 2000. MesP1 and MesP2 are essential for the development of cardiac mesoderm. Development, 127(15), pp.3215–3226.

Koch, F. et al., 2017. Antagonistic Activities of Sox2 and Brachyury Control the Fate Choice of Neuro-Mesodermal Progenitors. Developmental Cell, 42(5), pp.514–526.e7.

Lawson, K.A., 1999. Fate mapping the mouse embryo. The International Journal of Developmental Biology, 43(7), pp.773–775.

Lawson, K.A. and Pedersen, R.A., 1987. Cell fate, morphogenetic movement and population kinetics of embryonic endoderm at the time of germ layer formation in the mouse. Development, 101(3), pp.627–652.

Lawson, K.A., Meneses, J.J. and Pedersen, R.A., 1986. Cell fate and cell lineage in the endoderm of the presomite mouse embryo, studied with an intracellular tracer. Developmental Biology, 115(2), pp.325–339.

Lawson, K.A., Meneses, J.J. and Pedersen, R.A., 1991. Clonal analysis of epiblast fate during germ layer formation in the mouse embryo. Development, 113(3), pp.891–911.

Lescroart, F. et al., 2018. Defining the earliest step of cardiovascular lineage segregation by single-cell RNA-seq. Science, 359(6380), pp.1177–1181.

Lescroart, F. et al., 2014. Early lineage restriction in temporally distinct populations of Mesp1 progenitors during mammalian heart development. Nature Cell Biology.

Li, H. and Durbin, R., 2010. Fast and accurate long-read alignment with Burrows-Wheeler transform. Bioinformatics (Oxford, England), 26(5), pp.589–595.

Liu, P. et al., 1999. Requirement for Wnt3 in vertebrate axis formation. Nature Genetics, 22(4), pp.361–365.

Lun, A.T.L., McCarthy, D.J. and Marioni, J.C., 2016. A step-by-step workflow for low-level analysis of single-cell RNA-seq data with Bioconductor. F1000Research, 5, p.2122.

Mohammed, H. et al., 2017. Single-Cell Landscape of Transcriptional Heterogeneity and Cell Fate Decisions during Mouse Early Gastrulation. CellReports, 20(5), pp.1215–1228.

Monaghan, A.P. et al., 1993. Postimplantation expression patterns indicate a role for the mouse forkhead/HNF-3 alpha, beta and gamma genes in determination of the definitive endoderm, chordamesoderm and neuroectoderm. Development, 119(3), pp.567–578.

Muzumdar, M.D. et al., 2007. A global double-fluorescent Cre reporter mouse. genesis, 45(9), pp.593–605.

Nowotschin, S. et al., 2013. The T-box transcription factor Eomesodermin is essential for AVE induction in the mouse embryo. Genes and Development, 27(9), pp.997–1002.

Parameswaran, M. and Tam, P.P., 1995. Regionalisation of cell fate and morphogenetic movement of the mesoderm during mouse gastrulation. Developmental genetics, 17(1), pp.16–28.

Park, E.J. et al., 2008. System for tamoxifen-inducible expression of cre-recombinase from the Foxa2 locus in mice. Developmental Dynamics, 237(2), pp.447–453.

Peng, G. et al., 2019. Molecular architecture of lineage allocation and tissue organization in early mouse embryo. Nature, 572(7770), pp.528–532.

Pijuan-Sala, B. et al., 2019. A single-cell molecular map of mouse gastrulation and early organogenesis. Nature, 566(7745), pp.490–495.

Pimeisl, I.-M. et al., 2013. Generation and characterization of a tamoxifen-inducible Eomes CreERmouse line. genesis, 51(10), pp.725–733.

Probst, S. and Arnold, S.J., 2017. Eomesodermin-At Dawn of Cell Fate Decisions During Early Embryogenesis. Current topics in developmental biology, 122, pp.93–115.

Probst, S. et al., 2017. A dual-fluorescence reporter in the Eomes locus for live imaging and medium-term lineage tracing. genesis, 55(8), p.e23043.

Rivera-Pérez, J.A. and Hadjantonakis, A.-K., 2014. The Dynamics of Morphogenesis in the Early Mouse Embryo. Cold Spring Harbor Perspectives in Biology, 7(11), p.a015867.

Rivera-Pérez, J.A., Mager, J. and Magnuson, T., 2003. Dynamic morphogenetic events characterize the mouse visceral endoderm. Developmental Biology, 261(2), pp.470–487.

Robertson, E.J., 2014. Dose-dependent Nodal/Smad signals pattern the early mouse embryo. Seminars in cell and developmental biology, 32, pp.73–79.

Saga, Y. et al., 1999. MesP1 is expressed in the heart precursor cells and required for the formation of a single heart tube. Development, 126(15), pp.3437–3447.

Sako, K. et al., 2016. Optogenetic Control of Nodal Signaling Reveals a Temporal Pattern of Nodal Signaling Regulating Cell Fate Specification during Gastrulation. CellReports, 16(3), pp.866–877.

Sasaki, H. and Hogan, B.L., 1993. Differential expression of multiple fork head related genes during gastrulation and axial pattern formation in the mouse embryo. Development, 118(1), pp.47–59.

Scialdone, A. et al., 2016. Resolving early mesoderm diversification through single-cell expression profiling. Nature, 535(7611), pp.289–293.

Snow, M.H.L., 1977. Gastrulation in the mouse: Growth and regionalization of the epiblast. Development, 42(1), pp.293–303.

Stuart, T. et al., 2019. Comprehensive Integration of Single-Cell Data. Cell, 177(7), pp.1888–1902.e21.

Tam, P.P. and Beddington, R.S., 1987. The formation of mesodermal tissues in the mouse embryo during gastrulation and early organogenesis. Development, 99(1), pp.109–126.

Tam, P.P. and Behringer, R.R., 1997. Mouse gastrulation: the formation of a mammalian body plan. Mechanisms of Development, 68(1–2), pp.3–25.

Tam, P.P. et al., 1997. The allocation of epiblast cells to the embryonic heart and other mesodermal lineages: the role of ingression and tissue movement during gastrulation. Development, 124(9), pp.1631–1642.

Tam, P.P.L. et al., 2007. Sequential allocation and global pattern of movement of the definitive endoderm in the mouse embryo during gastrulation. Development, 134(2), pp.251–260.

Teo, A.K.K. et al., 2011. Pluripotency factors regulate definitive endoderm specification through eomesodermin. Genes and Development, 25(3), pp.238–250.

Tosic, J. et al., 2019. Eomes and Brachyury control pluripotency exit and germ-layer segregation by changing the chromatin state. Nature Cell Biology, 10, pp.91–14.

Tzouanacou, E. et al., 2009. Redefining the Progression of Lineage Segregations during Mammalian Embryogenesis by Clonal Analysis. Developmental Cell, 17(3), pp.365–376.

van Boxtel, A.L. et al., 2015. A Temporal Window for Signal Activation Dictates the Dimensions of a Nodal Signaling Domain. Developmental Cell, 35(2), pp.175–185.

van Boxtel, A.L. et al., 2018. Long-Range Signaling Activation and Local Inhibition Separate the Mesoderm and Endoderm Lineages. Developmental Cell, 44(2), pp.179–191.e5.

van den Ameele, J. et al., 2012. Eomesodermin induces Mesp1 expression and cardiac differentiation from embryonic stem cells in the absence of Activin. EMBO reports, 13(4), pp.355–362.

Vincent, S.D. and Robertson, E.J., 2003. Highly efficient transgene-independent recombination directed by a maternally derived SOX2CRE transgene. genesis, 37(2), pp.54–56.

Vincent, S.D. et al., 2003. Cell fate decisions within the mouse organizer are governed by graded Nodal signals. Genes and Development, 17(13), pp.1646–1662.

Viotti, M., Foley, A.C. and Hadjantonakis, A.-K., 2014. Gutsy moves in mice: cellular and molecular dynamics of endoderm morphogenesis. Philosophical transactions of the Royal Society of London. Series B, Biological sciences, 369(1657), pp.20130547–20130547.

Viotti, M., Nowotschin, S. and Hadjantonakis, A.-K., 2014. SOX17 links gut endoderm morphogenesis and germ layer segregation. Nature Cell Biology, 16(12), pp.1146–1156.

Wang, Q. et al., 2017. The p53 Family Coordinates Wnt and Nodal Inputs in Mesendodermal Differentiation of Embryonic Stem Cells. Cell Stem Cell, 20(1), pp.70–86.

Weinstein, D.C. et al., 1994. The winged-helix transcription factor HNF-3 beta is required for notochord development in the mouse embryo. Cell, 78(4), pp.575–588.

Wen, J. et al., 2017. Single-cell analysis reveals lineage segregation in early post-implantation mouse embryos. The Journal of biological chemistry, 292(23), pp.9840–9854.

Wymeersch, F.J. et al., 2016. Position-dependent plasticity of distinct progenitor types in the primitive streak. eLife, 5, p.e10042.

