## Supplemental_information for "Spatiotemporal sequence of mesoderm and endoderm lineage segregation during mouse gastrulation"

Figure S1

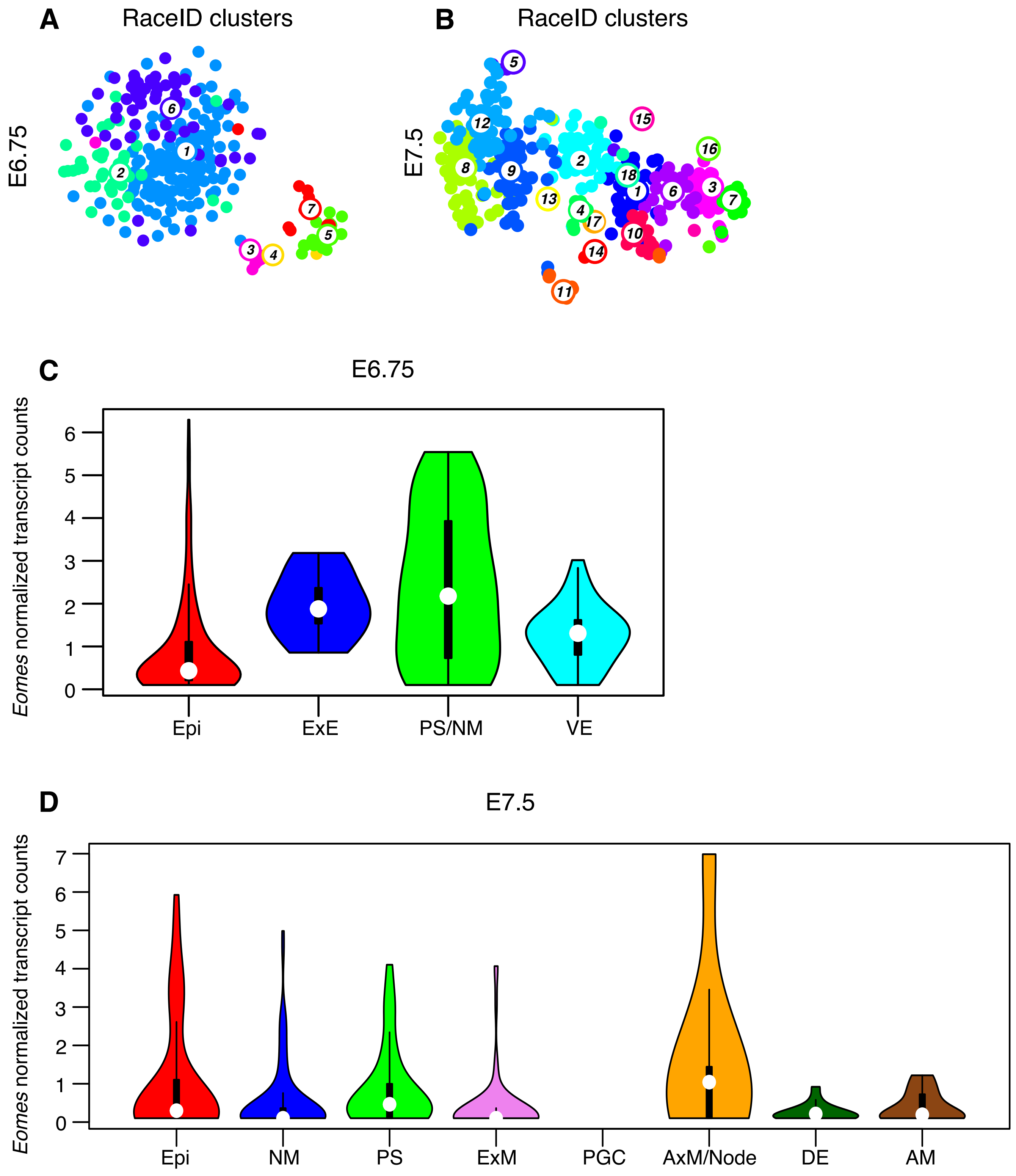


**Figure S1: RaceID3 clusters and expression levels of *Eomes* in different cell populations**

**A**, **B**) t-SNE plots showing clusters identified by the RaceID3 algorithm in single cells sequenced from E6.75 (**A**) and E7.5 (**B**) embryos. **C**, **D**) Violin plots showing *Eomes* expression by normalized transcript counts in assigned clusters from E6.75 embryos (**C**) and E7.5 embryo (**D**). Anterior mesoderm (AM), axial mesoderm (AxM), definitive endoderm (DE), epiblast (Epi), nascent mesoderm (NM), extraembryonic ectoderm (ExE), extraembryonic mesoderm (ExM), primordial germ cell (PGC), primitive streak (PS), visceral endoderm (VE).

Figure S2

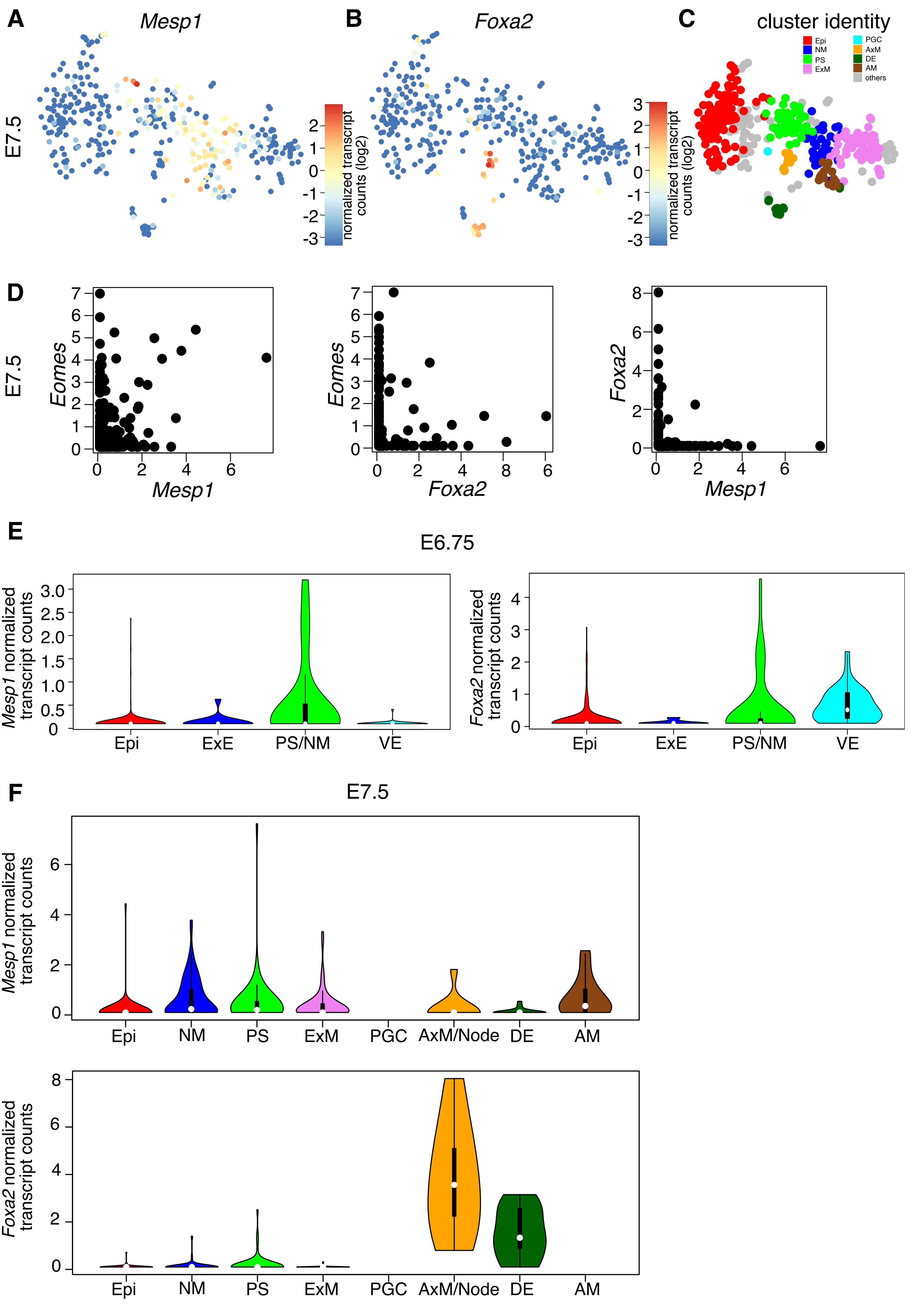
A
Mesp1
B
Foxa2
C
cluster identity
E7.5
normalized transcript counts (log2)
Epi
NM
PS
ExM
PGC
AxM
DE
AM
others
D
Eomes
Mesp1
Foxa2
E
E6.75
Mesp1 normalized transcript counts
Foxa2 normalized transcript counts
Epi
ExE
PS/NM
VE
F
E7.5
Epi
NM
PS
ExM
PGC
AxM/Node
DE
AM

**Figure S2: Spatial separation of *Mesp1* and *Foxa2* positive populations at E7.5**
**A**, **B**) t-SNE maps of E7.5 scRNA-seq data showing *Mesp1*- (**A**) and *Foxa2*- (**B**) expressing cells. The scale represents log2 normalized transcript counts. **C**) t-SNE plot with assigned identities to different clusters at E7.5. **D**) Scatter plots of single cells from E7.5 timepoint for *Eomes/Mesp1*, *Eomes/Foxa2* and *Foxa2/Mesp1* expression. Only one *Foxa2/Mesp1* double positive cell is detected. X- and y-axes indicate normalized transcript counts. **E**, **F**) Violin plots showing *Mesp1* and *Foxa2* expression by normalized transcript counts in in assigned clusters from E6.75 embryos (**E**) and E7.5 embryos (**F**). Anterior mesoderm (AM), axial mesoderm (AxM), definitive endoderm (DE), epiblast (Epi), nascent mesoderm (NM), extraembryonic ectoderm (ExE), extraembryonic mesoderm (ExM), primordial germ cell (PGC), primitive streak (PS), visceral endoderm (VE).

Figure S3

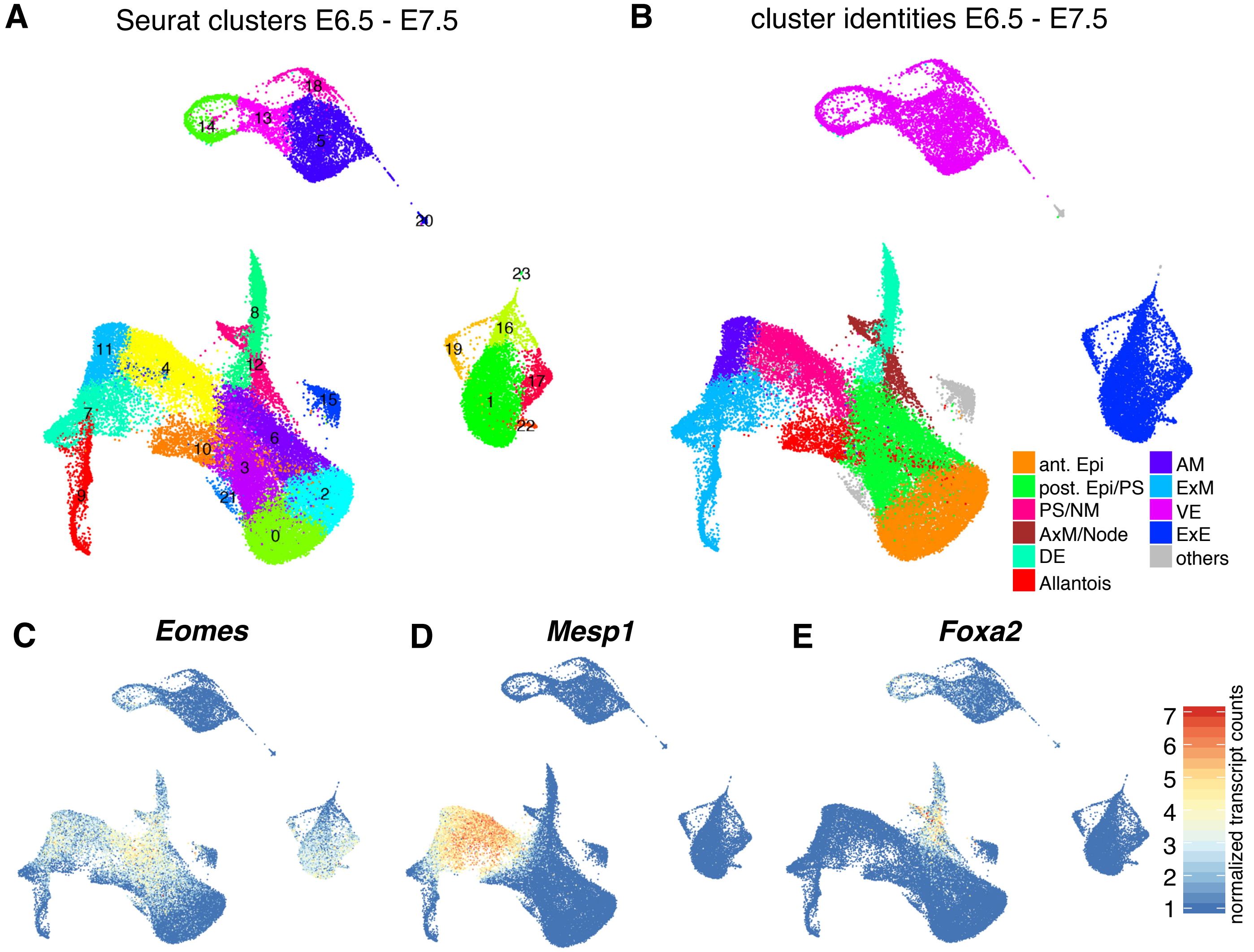


F

| Timepoint | Total # of cells | Double+ cells | *Mesp1*+ cells | *Foxa2*+ cells | Total # *Mesp1*+ and *Foxa2*+ cells | % double + cells |
|---|---|---|---|---|---|---|
| Pijuan-Sala et al 2019 data | | | | | | |
| 6.5 | 1000 | 1 | 24 | 14 | 37 | 2,70% |
| 6.75 | 1364 | 11 | 230 | 105 | 324 | 3,40% |
| 7.0 | 11048 | 211 | 2865 | 1604 | 4258 | 5,00% |
| 7.25 | 10317 | 179 | 3516 | 1274 | 4611 | 3,90% |
| 7.5 | 7196 | 54 | 2204 | 490 | 2640 | 2,00% |
| our data | | | | | | |
| 6.75 | 239 | 1 | 23 | 37 | 59 | 1,70% |
| 7.5 | 371 | 3 | 101 | 36 | 134 | 2,20% |

**Figure S3: Analysis of a published scRNA-seq dataset with large cell numbers**
**A**, **B**) UMAP representations showing clusters identified with the Seurat package (**A**) and assigned identities to different clusters (**B**) of the timepoints E6.5, E6.75, E7.0, E7.25, and E7.5 combined (data from (Pijuan-Sala et al. 2019)). Anterior mesoderm (AM), axial mesoderm (AxM) definitive endoderm (DE), anterior/posterior epiblast (ant./post. Epi), nascent mesoderm (NM), extraembryonic ectoderm (ExE), extraembryonic mesoderm (ExM), primitive streak (PS), visceral endoderm (VE). **C**-**E**) UMAP representations showing the expression of *Eomes* (**C**), *Mesp1* (**D**) and *Foxa2* (**E**). The scale represents normalized transcript counts. **F**) Table showing the numbers of *Mesp1* positive, *Foxa2* positive, and *Mesp1/Foxa2* double positive cells at the different timepoints analyzed by (Pijuan-Sala et al., 2019) and in this paper. The percentage of double positive cells within the *Mesp1* and *Foxa2* positive cells is indicated.

Figure S4

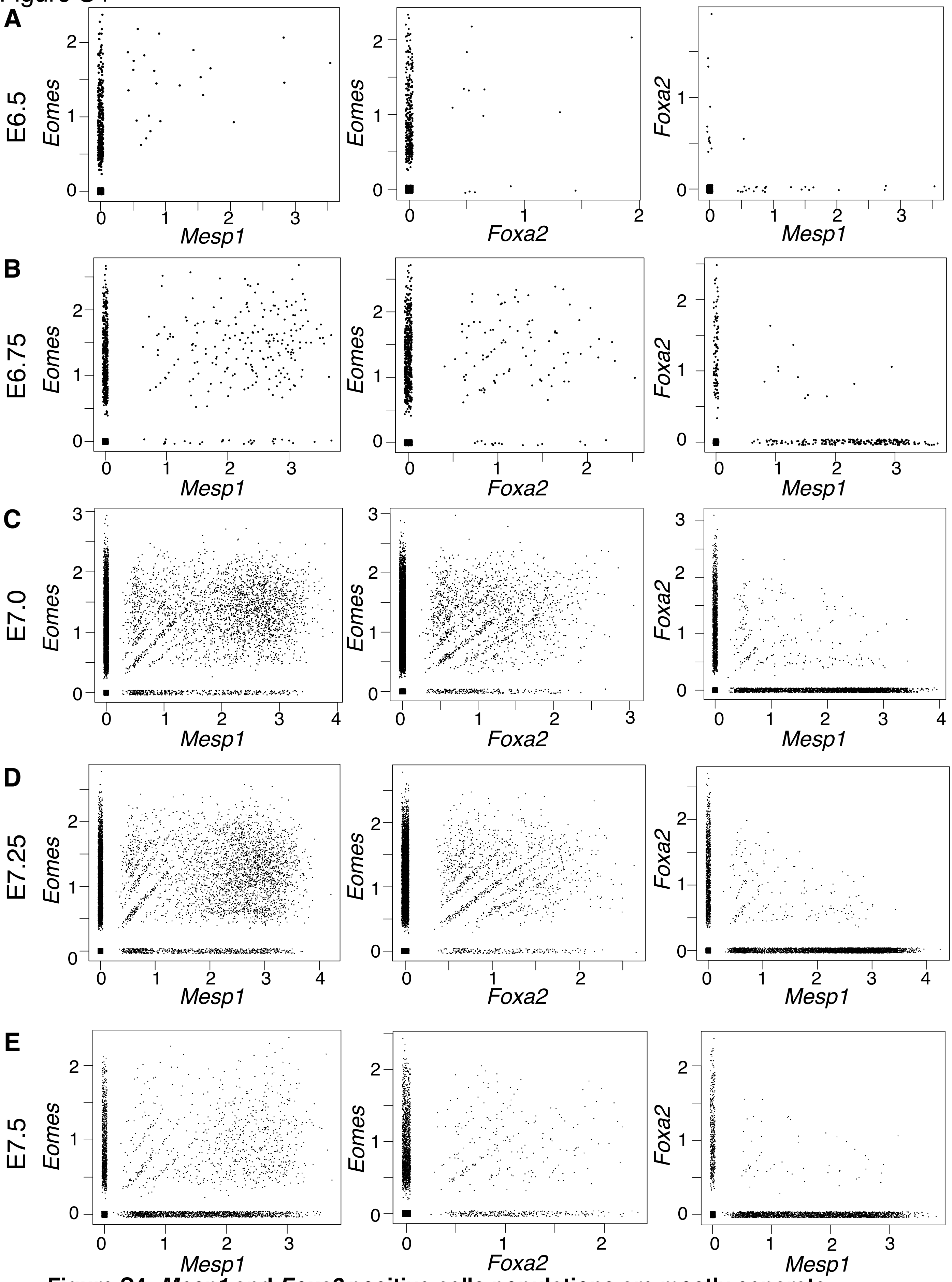


**Figure S4: *Mesp1* and *Foxa2* positive cells populations are mostly separate**
**A-E**) Scatter plots of single cells from separate timepoints for *Eomes/Mesp1*, *Eomes/Foxa2* and *Foxa2/Mesp1* expression. *Foxa2/Mesp1* double positive constitute a minor population (data from (Pijuan-Sala et al., 2019). X- and y-axes indicate normalized transcript counts. (**A**) E6.5, (**B**) E6.75, (**C**) E7.0, (**D**) E7.25, and (**E**) E7.5.

Figure S5

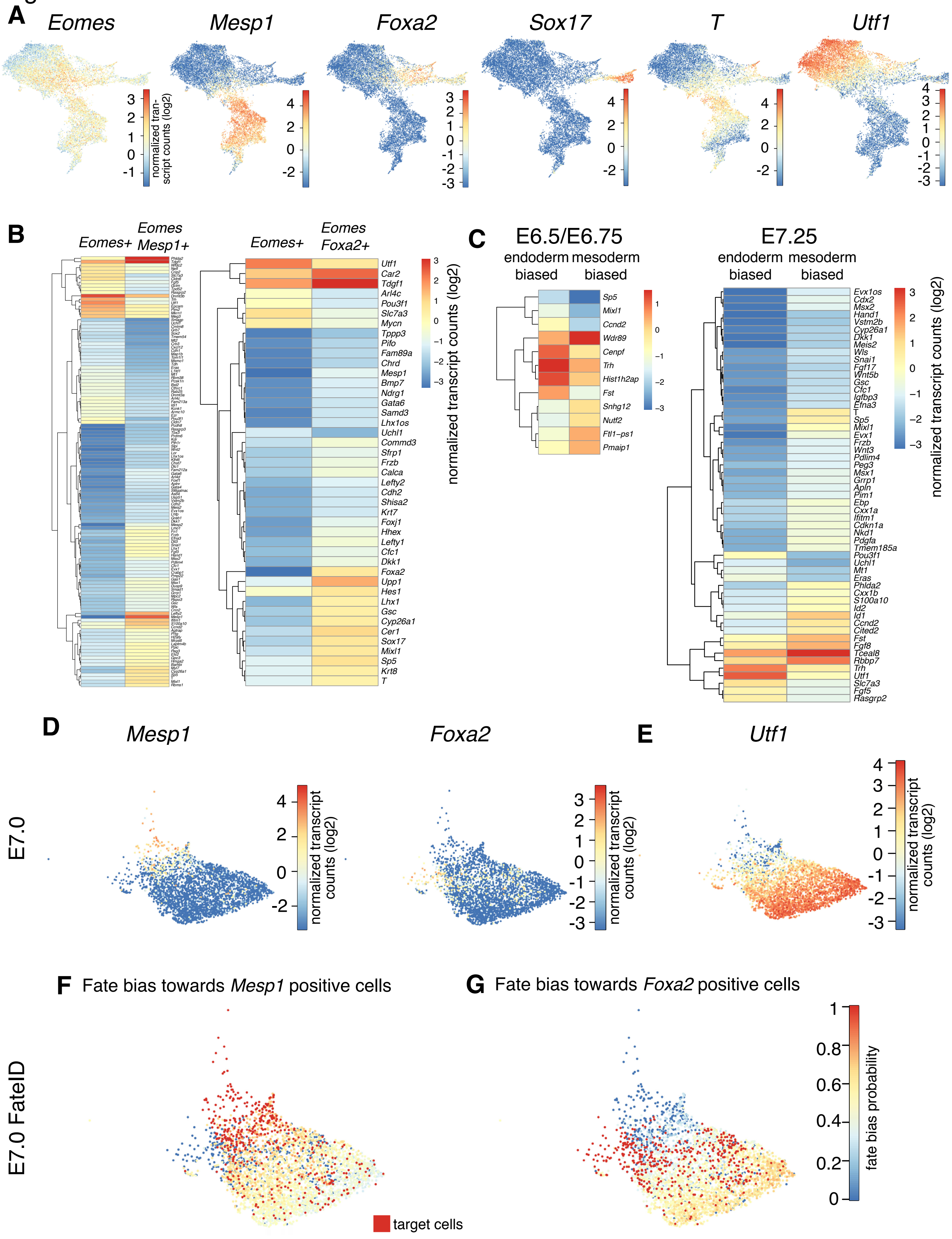
A
Eomes
Mesp1
Foxa2
Sox17
T
Utf1
normalized tran- script counts (log2)
B
Eomes+
Eomes Mesp1+
Eomes+
Eomes Foxa2+
Utf1
Car2
Tdgf1
Arl4c
Pou3f1
Slc7a3
Mycn
Tppp3
Pifo
Fam89a
Chrd
Mesp1
Bmp7
Ndrg1
Gata6
Samd3
Lhx1os
Uchl1
Commd3
Sfrp1
Frzb
Calca
Lefty2
Cdh2
Shisa2
Krt7
Foxj1
Hhex
Lefty1
Cfc1
Dkk1
Foxa2
Upp1
Hes1
Lhx1
Gsc
Cyp26a1
Cer1
Sox17
Mixl1
Sp5
Krt8
T
normalized transcript counts (log2)
C
E6.5/E6.75
endoderm biased
mesoderm biased
Sp5
Mixl1
Ccnd2
Wdr89
Cenpf
Trh
Hist1h2ap
Fst
Snhg12
Nutf2
Ftl1-ps1
Pmaip1
E7.25
endoderm biased
mesoderm biased
Evx1os
Cdx2
Msx2
Hand1
Vstm2b
Cyp26a1
Dkk1
Meis2
Wls
Snai1
Fgf17
Wnt5b
Gsc
Cfc1
Igfbp3
Efna3
T
Sp5
Mixl1
Evx1
Frzb
Wnt3
Pdlim4
Peg3
Msx1
Grrp1
Apln
Pim1
Ebp
Cxx1a
Ifitm1
Cdkn1a
Nkd1
Pdgfa
Tmem185a
Pou3f1
Uchl1
Mt1
Eras
Phlda2
Cxx1b
S100a10
Id2
Id1
Ccnd2
Cited2
Fst
Fgf8
Tceal8
Rbbp7
Trh
Utf1
Slc7a3
Fgf5
Rasgrp2
normalized transcript counts (log2)
D
E7.0
Mesp1
Foxa2
normalized transcript counts (log2)
E
Utf1
normalized transcript counts (log2)
F
Fate bias towards Mesp1 positive cells
G
Fate bias towards Foxa2 positive cells
E7.0 FateID
target cells
fate bias probability

**Figures S5: Fate bias analysis in *Eomes* single positive posterior epiblast cells**

**A**) UMAP representations of embryonic *Eomes* positive cells from E6.5 to E7.5 with an expression cut off of >0.3 normalized transcript counts showing levels of *Eomes, Mesp1, Foxa2, Sox17, T,* and *Utf1* expression. **B**) Heat map representation of differentially expressed genes (>2-fold) between *Eomes* single positive cells and *Eomes/Foxa2* double positive cells or *Eomes/Mesp1* double positive cells. Scale bar represents log2 normalized transcript counts. **C**) Heat map representation of differentially expressed genes (>2-fold) between *Eomes* single positive cells biased towards *Foxa2* positive cells (endoderm biased) and *Eomes* single positive cells biased towards *Mesp1* positive cells (mesoderm biased) at timepoints E6.5/E6.75 and E7.25. Scale bar represents log2 normalized transcript counts. **D**, **E**) *Eomes* positive cells from E7.0 were clustered and *Mesp1* and *Foxa2* (**D**), and *Utf1* (**E**) expression was plotted onto the UMAP representation. Scale bar represents log2 normalized transcript counts. **F**, **G**) FateID analysis of embryonic *Eomes* positive cells from timepoint E7.0. The fate bias probability is indicated in single *Eomes* positive cells towards *Mesp1* positive cells (**F**) and *Foxa2* positive cells (**G**) (red, target cells). Color scale represents fate bias probabilities on the scale from 0 to 1. Data from (Pijuan-Sala et al., 2019).

**Supplementary Table Legends**

**Supplementary Table 1**
Differentially expressed genes in each of the seven identified clusters compared to all other clusters in the scRNA-seq data at E6.75.

**Supplementary Table 2**
Differentially expressed genes in each of the 18 identified clusters compared to all other clusters in the scRNA-seq data at E7.5.

**Supplementary Table 3**
Counts table of scRNA-seq data at E6.75, including expression data of handpicked marker genes in *Eomes* single positive, *Eomes/Mesp1* double positive, and *Eomes/Foxa2* double positive cells. Lanes colored in light red on the second sheet are from extraembryonic clusters.

**Supplementary Table 4**
Counts table of scRNA-seq data at E7.5, including expression data of handpicked marker genes in *Eomes* single positive, *Eomes/Mesp1* double positive, and *Eomes/Foxa2* double positive cells.

**Supplementary Table 5**
Differentially expressed genes in *Eomes/Mesp1* or *Eomes/Foxa2* double positive cells compared to *Eomes* single positive cells.

**Supplementary Table 6**
Differentially expressed genes between endoderm and mesoderm biased *Eomes* single positive cells at E6.5/E6.75, E7.0, and E7.25.
